# Phosphoglycerate Kinase Can Adopt a Topologically Misfolded Form that is More Stable than its Native State

**DOI:** 10.1101/2025.06.24.661412

**Authors:** Yingzi Xia, Barbara Amann, Richard E. Gillilan, Piyoosh Sharma, Sreemantee Sen, Karen G. Fleming, Stephen D. Fried

## Abstract

The native states of globular proteins are typically viewed as being the most stable conformations on their respective proteins’ soluble free energy landscapes. This view, known as the Thermodynamic Hypothesis, explains why many proteins can reversibly refold after being denatured. Here we report an intriguing counterexample to this paradigm. When *E. coli* phosphoglycerate kinase (PGK) is stimulated to refold upon dilution from denaturant, instead of returning to its native state, it populates an unusual misfolded form that is monomeric and native-like, but which is even more kinetically stable than its native form, as based on its resistance to thermal and detergent-induced denaturation. Moreover, this misfolded form cannot self-correct, even for days. We show that the key structural feature of this misfolded form of PGK is topological in nature by demonstrating that kinetically stable misfolded forms do not form any longer if PGK is circularized, which prevents its termini from threading through other portions of the protein. Our findings demonstrate that a misfolded protein need not aggregate or form an amyloid to become stabilized with respect to the native state, and call attention to topologically-misfolded proteins as a potential Achilles heel to the cellular proteostasis network.

## Introduction

According to the Thermodynamic Hypothesis, native states of proteins represent global thermodynamic minima and are the conformations that minimize their Gibbs free energy under physiological conditions (*1–4*). This theory provides a framework for why many small globular proteins can efficiently refold spontaneously after being fully denatured (*5–7*). On the other hand, many proteins can occupy metastable intermediates or misfolded forms. Amyloid fibrils represent a special case of misfolding which possess highly stabilized cross-beta structures (*8–11*); because of this stability, amyloids pose unique challenges to the proteostasis network, though some disaggregase chaperones possess the capacity to solubilize them (*12–17*). On the other hand, it is generally thought that with amyloids and other aggregates states excluded, the Thermodynamic Hypothesis still prevails, and that native states represent global minima on the soluble portion of proteins’ energy landscapes (*18*). This model explains why soluble misfolded states can often be resolved by molecular chaperones (*18–21*), which through a combination of foldase and unfoldase activities, facilitate thermodynamically-driven transitions from metastable misfolded forms to the native state.

In previous work, we estimated that ca. 40% of *E. coli*’s soluble proteome cannot spontaneously refold under typical *in vitro* refolding conditions (overnight incubation in 6 M guanidinium chloride (GdmCl), followed by dilution to 0.06 M GdmCl in 20 mM Tris, pH 8.0) (*22*), a fraction which decreases to 15-20% when chaperones like GroEL (Hsp60) or DnaK (Hsp70) are supplied (*23*). 105 proteins still did not efficiently refold even with chaperone assistance, and yet this smaller group still included the majority of *E. coli*’s glycolytic enzymes (8 out of 9). Phosphoglycerate kinase (PGK) is one such protein. We were curious to interrogate why it cannot refold following denaturation – What types of misfolded states does it form and what makes them inefficient at self-correcting or getting recognized by molecular chaperones?

Over the years, PGK has been used as a model to study the folding of a two-domain protein (*24–28*). The N-terminal subdomain binds 3-phosphoglycerate and the C-terminal domain binds ATP; closure between these two domains enables the enzyme to catalyze a phosphoryl group transfer between the two substrates (*29*). In eukaryotic orthologues of PGK (from horse and yeast), the two domains fold independently of one another and the full-length protein refolds with a metastable intermediate (*30, 31*). On the other hand, *E.* coli PGK is 5 orders of magnitude more kinetically stable than yeast PGK and its C-terminal domain cannot fold independently of the N-terminal domain (*32*). Non-reversible behaviour has been previously observed in its capacity to regain activity after being chemically denatured (*27, 33*). In yeast PGK, unusually compact non-native intermediates have been detected by single-molecule FRET (*34*), and stretched exponential kinetics have been observed by fluorescence (*35, 36*).

More recently, we found by combining simulations and structural mass spectrometry that upon attempting to refold from a fully unfolded state, PGK (from both *E. coli* and yeast) occupies several misfolded states that possess changes in entanglement status – either a gain in entanglement absent in the native state, or a loss of entanglement present in the native state (*37*). These misfolded structures specifically entail non-covalent lasso entanglements (*38, 39*), a topological change involving two interacting residues forming a “loop” through which another portion of the protein (the threading segment) passes through (*40–44*). We showed these entangled misfolded forms explain the origin of stretched exponential kinetics; however, experimental evidence that they provide kinetic traps deep enough to make refolding to the native state irreversible is lacking. Moreover, there is currently very little direct experimental characterization of entangled misfolded states in general. How a non-native entanglement might impact structure, activity, and stability has not yet been demonstrated experimentally. And unambiguous experimental evidence for their existence is still lacking.

Here, we report that when *E. coli* PGK is refolded by dilution following an extended incubation in denaturant, it populates conformation(s) that are soluble, monomeric, and with native-like secondary structure, but which have low enzymatic activity. These misfolded conformations also possess unusually high kinetic stability (*45*), as evidenced by high resistance to thermal denaturation and detergent. To support the view that this unusual set of features is due to a topological change, we show that when PGK is circularized (which precludes the termini from threading through loops), it no longer populates kinetically-stable misfolded states. Curiously, we find that PGK can refold efficiently if it is denatured for shorter durations; entangled misfolded forms *only* populate after diluting out of denaturant following an extended exposure to it. Hence, we show that PGK’s equilibrated unfolded ensemble possesses structure that seeds the formation of entanglements. In summary, we provide experimental evidence for the existence of soluble topologically misfolded states, which exhibit impaired activity but unusually high stability. The potential challenges this mode of misfolding mounts to the proteostasis network (*46, 47*) and its implications to protein folding disease is discussed.

## Results

Wild-type PGK was purified from *E. coli* (Figure S1A-C) and chemically refolded by first incubating in 6.0 M guanidinium chloride (GmdCl) for either 2 or 24 h (to form U2, U24), and then diluting 100-fold (Figure 1A). We first confirmed through a GdmCl titration that PGK’s midpoint concentration is 0.63 ± 0.12 M GdmCl, implying that in 6 M GdmCl, PGK is unfolded, and that 0.06 M GdmCl falls well within the native baseline (Figure S1D). Moreover, the thermodynamic stability of *E. coli* PGK found from fitting the titration was within error to a previously reported value found by denaturing with urea (Figure S1D) (*32*). As much of our study focuses on comparing natively expressed PGK to a chemically refolded form, we developed a procedure to ensure that native and refolded PGK samples are consistently prepared in identical buffers with identical concentrations, so that samples differ only in history but are compositionally identical (Figure S1A, see Methods). Far-UV circular dichroism (CD) spectra showed that after 1 h of refolding, PGK could reform its native structure if was initially unfolded for 2 h (cf. spectra of native (N) and U2-R1, Figure 1B; all raw CD data compiled in Data S1). However, if PGK is first unfolded for 24 h and then allowed 1 h to refold (U24-R1), it produces a consistently altered CD spectrum (Figure 1B-C) with less dichroism between 220-230 nm, suggesting the formation of a near-native misfolded state. With analytical ultracentrifugation (AUC), we confirmed that the chemically refolded forms of PGK are soluble and monomeric with identical sedimentation velocities compared to the native enzyme (Figure 1D; all fitted sedimentation coefficients are compiled in Table S1 and raw AUC data in Data S2). However, unlike native PGK and U2-R1, which have apparent melting temperatures at 60 °C based on CD thermal melts, U24-R1 does not thermally denature upto 95°C (Figure 1E-F). Hence, U24-R1 PGK populates a set of conformations that are more kinetically stable with respect to temperature than native PGK. Further CD spectroscopy experiments showed that native PGK irreversibly thermally denatures at 95°C (Figure S2A-B), whilst U24-R1 PGK possesses folded structure when incubated upto 95°C that is retained upon cooling (Figure S2C-D). If incubated at 95°C for 24 h, U24-R1 eventually melts (Figure S2E), confirming that the effect we observed is kinetic in nature (as thermodynamics would require; N and U24-R1 are after all the same molecule in the same conditions).

**Figure 1.**
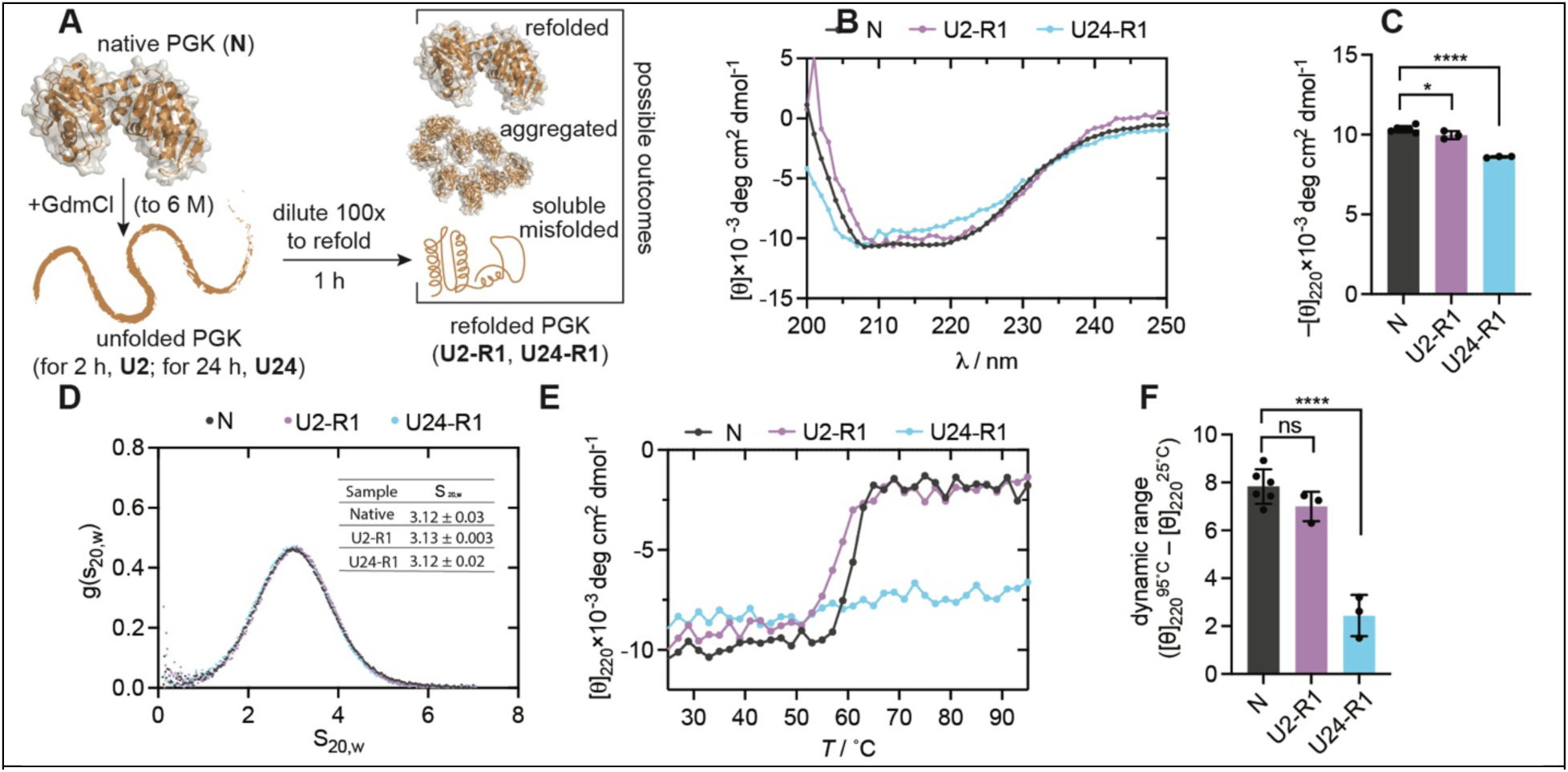
PGK Misfolding Produces a Soluble Kinetically Stable State. (A) Scheme illustrating how PGK is chemically unfolded and refolded and nomenclature for unfolded states based on unfolding time (U2, U24) and their corresponding refolded states (U2-R1, U24-R1). (B) Representative far-UV circular dichroism (CD) spectra of native (N) and chemically unfolded-refolded forms of PGK. (C) Technical replicate (*n* ≥ 3) values for mean residual ellipticity (MRE) of forms of PGK at 220 nm. P-values calculated by one-way ANOVA with Bonferroni’s multiple comparison test. *, P< 0.05; ****, P < 0.0001. Error bars are std. devs. (D) Representative sedimentation velocity distributions for native and chemically unfolded-refolded forms of PGK. Inset table shows mean ± std. dev. for sedimentation velocity normalized to 20°C in pure water (S_20,w_) from technical replicates (*n* ≥ 2). (E) Representative thermal melts of PGK as monitored by CD signal at 220 nm either on native enzyme or on chemically unfolded-refolded forms, ramping at 1 °C/min upto 95°C. (F) Difference in MRE at 95°C endpoint and 25°C starting point for technical replicates (*n* ≥ 3) of the thermal melt. P-values calculated by one-way ANOVA with Bonferroni’s multiple comparison test. ns, not significant; ****, P < 0.0001. Error bars are std. devs.

Stimulated by the finding that U24-R1 possesses greater apparent thermal stability than native PGK, we sought an alternative assay to assess its kinetic stability and performed detergent resistance assays (Figure 2). In this experiment, PGK is incubated in 0.6% SDS, incubated at 25°C or 90°C, and resolved by denaturing SDS-PAGE. Whilst most proteins denature in 0.6% SDS, highly kinetically stable proteins can resist detergent denaturation (*48, 49*). On the other hand, boiling in 0.6% SDS unfolds even exceptionally stable proteins. We find that native PGK and U2-R1 are denatured when incubated in SDS and migrate as unfolded proteins regardless of whether they are boiled or not (Figure 2). However, U24-R1 resists SDS-induced denaturation and migrates with an unusual pattern (possibly corresponding to different conformations with distinct electrophoretic mobilities), unless it is boiled, at which point it unfolds and migrates like the other forms of PGK. This finding shows that a portion of the U24-R1 ensemble populates conformation(s) with high kinetic stability with respect to thermal and detergent denaturation.

**Figure 2.**
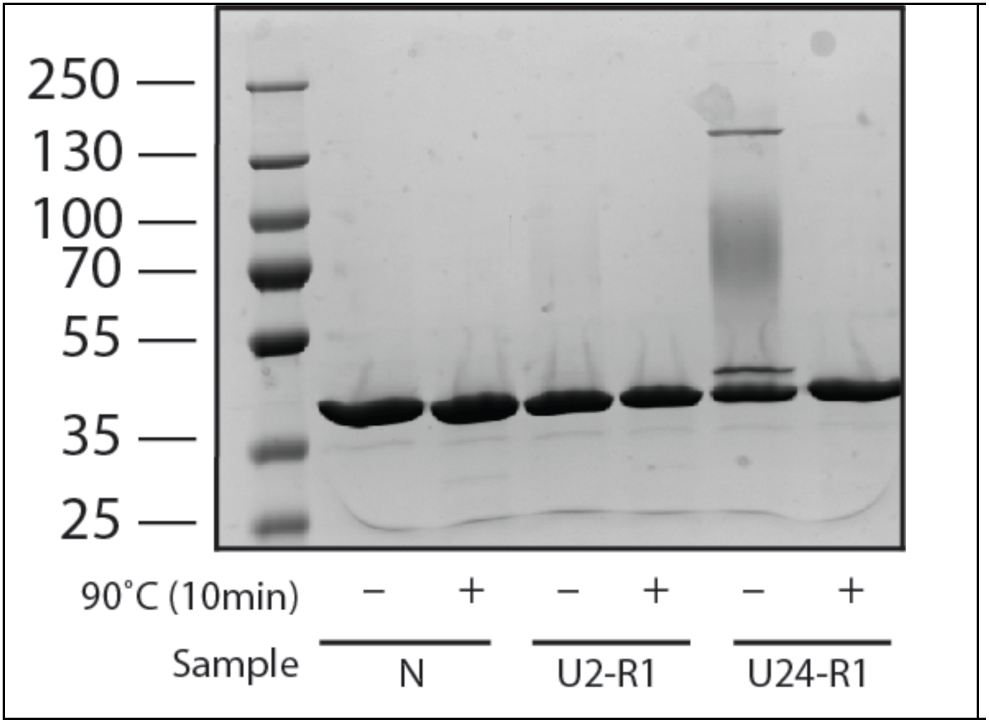
Detergent Resistance Assay of Misfolded PGK. Native (N) and chemically unfolded-refolded forms of PGK (U2-R1, U24-R1) were incubated in 0.6% SDS, boiled (or not), resolved by Tris-Glycine SDS-PAGE, and visualized with Coomassie stain. U24-R1 adopts conformations resistant to SDS denaturation.

PGK is a glycolytic enzyme that generates ATP by transferring phosphate from 1,3-bisphosphoglycerate to ADP (*50, 51*). Its activity is facile to measure by coupling the reverse reaction (ATP-dependent phosphorylation of 3-phosphoglycerate) to NADH depletion (Figure S3). Native PGK follows Michaelis-Menten kinetics with a *k*_cat_ of 135 ± 10 s^–1^ and *K*_M_ of 0.84 ± 0.09 mM (Figure 3A, D; all initial rates compiled in Data S3). In consonance with CD spectra, U2-R1 appears to be a *bona fide* refolded PGK with a *k*_cat_ that is identical to the native enzyme within error (Figure 3B, D). On the other hand, U24-R1 is significantly less active, with a *k*_cat_ of 10 ± 3 s^–1^ (14-fold lower than native; Figure 3C, D), albeit still with a detectable activity above background (Figure S3C). We repeated these Michaelis-Menten activity assays under which PGK was first unfolded in 8 M urea or under reducing conditions (6 M GdmCl and 10 mM dithiothreitol (DTT)). The nature of the unfolding conditions did not change the outcome: In all treatments, U2-R1 possessed a *k*_cat_ similar to that of native PGK, whilst U24-R1 possessed a *k*_cat_ ∼14-fold lower than native (Figure 3E-F). Hence, we find that U24-R1 populates a “near-native” misfolded form of PGK, which is soluble, monomeric and with similar secondary structure content, but nevertheless possesses low activity and unusually high stability.

**Figure 3.**
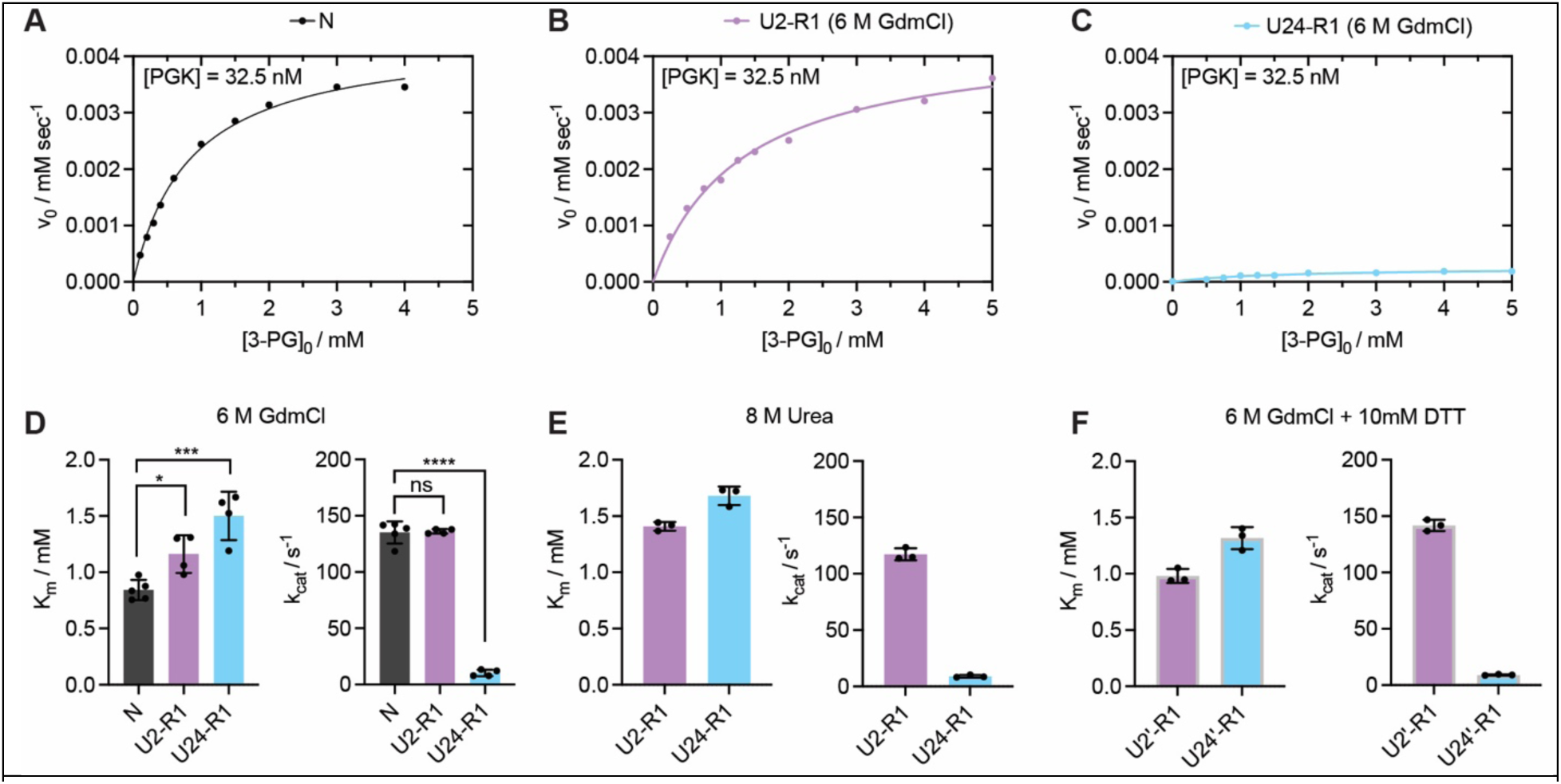
Enzymatic Characterization of Native and Chemically Unfolded-Refolded PGK. (A, B, C) Representative Michaelis-Menten curves showing initial rates as a function of initial substate concentration (3-phosphoglycerate) for (A) native PGK; (B) PGK unfolded for 2 h in 6 M GdmCl, diluted 100-fold and incubated for 1 h; (C) PGK unfolded for 24 h in 6 M GdmCl, diluted 100-fold and incubated for 1 h. (D) Michaelis constant (*K*_M_) and unimolecular rate constant (*k*_cat_) derived from fitting replicates (*n* ≥ 4) of datasets such as the one shown in panels A-C. Each dot represents the optimal value for *K*_M_ and *k*_cat_ based on a non-linear fit of the initial rates at 8-10 substrate concentrations. P-values calculated by one-way ANOVA with Bonferroni’s multiple comparison test. ns, not significant; *, P< 0.05; ***, P<0.001; ****, P < 0.0001. Error bars are std. devs. (E) *K*_M_ and *k*_cat_ derived from fitting replicates (*n* = 3) of Michaelis-Menten kinetics on chemically unfolded-refolded forms of PGK, except where PGK was unfolded in 8 M urea instead. Error bars are std. devs. (F) *K*_M_ and *k*_cat_ derived from fitting replicates (*n* = 3) of Michaelis-Menten kinetics on chemically unfolded-refolded forms of PGK, except where PGK was unfolded with 10 mM dithiothreitol (DTT) in addition to 6 M GdmCl. Error bars are std. devs.

To try to rationalize the structural basis for this unusual stability, we considered the possibility that U24-R1 populates entangled misfolded states, which either gain entanglements *not* present in the native structure, or lose entanglements that *are* present in the native structure. A recent growing body of evidence, primarily driven by computational modeling, suggests that non-covalent lasso entanglements are a pervasive feature in protein structure. They exist in many native protein structures (including PGK’s (*39, 44*)), and they can also be a key feature in misfolded conformations (*37, 52*). It has been suggested that entangled misfolded states may be quite stable in proteins with ≥300 amino acids because in order to resolve them, native portions of a protein must first unfold to allow threading segments to backtrack (*53*).

It has been challenging to obtain direct experimental evidence for proteins misfolding in a topological manner. However, we reasoned that such a form of misfolding might be dependent on the protein possessing free N- and C-termini capable of threading through loops closed by other native contacts in the protein. To test if the biophysical features of U24-R1 are dependent on it possessing free termini, we prepared a circularized version of PGK in which the N- and C-termini are fused together (Figure 4A), which was possible because luckily, PGK’s N- and C-termini are very proximal to each other in the protein’s native structure. We created PGK_circ_ through a sortase-mediated ligation (*54–56*), and purified it to homogeneity (Figure S4A) as based on SDS-PAGE and intact mass spectrometry (Figure S4B-E). Its CD spectrum is qualitatively similar to that of wild-type PGK (Figure S4F), though unlike wild-type PGK, PGK_circ_’s CD spectrum after unfolding-refolding (i.e., U24-R1) superimposes on top of its native spectrum. We reasoned that in such a topologically restrained state, PGK_circ_ might be restricted from misfolding in some of the ways wild-type PGK can (Figure 4B). Indeed, we found that after denaturing for 24 h and refolding for 1 h, unlike wild-type PGK, U24-R1 PGK_circ_ did not possess unusual resistance to SDS or thermal denaturation (Figure 4C-F). Circularizing PGK did however moderately boost the thermal stability of *both* the native and U24-R1 forms (Figure 4C), as expected (*57, 58*), given this new linkage will reduce the entropy of the unfolded state.

**Figure 4.**
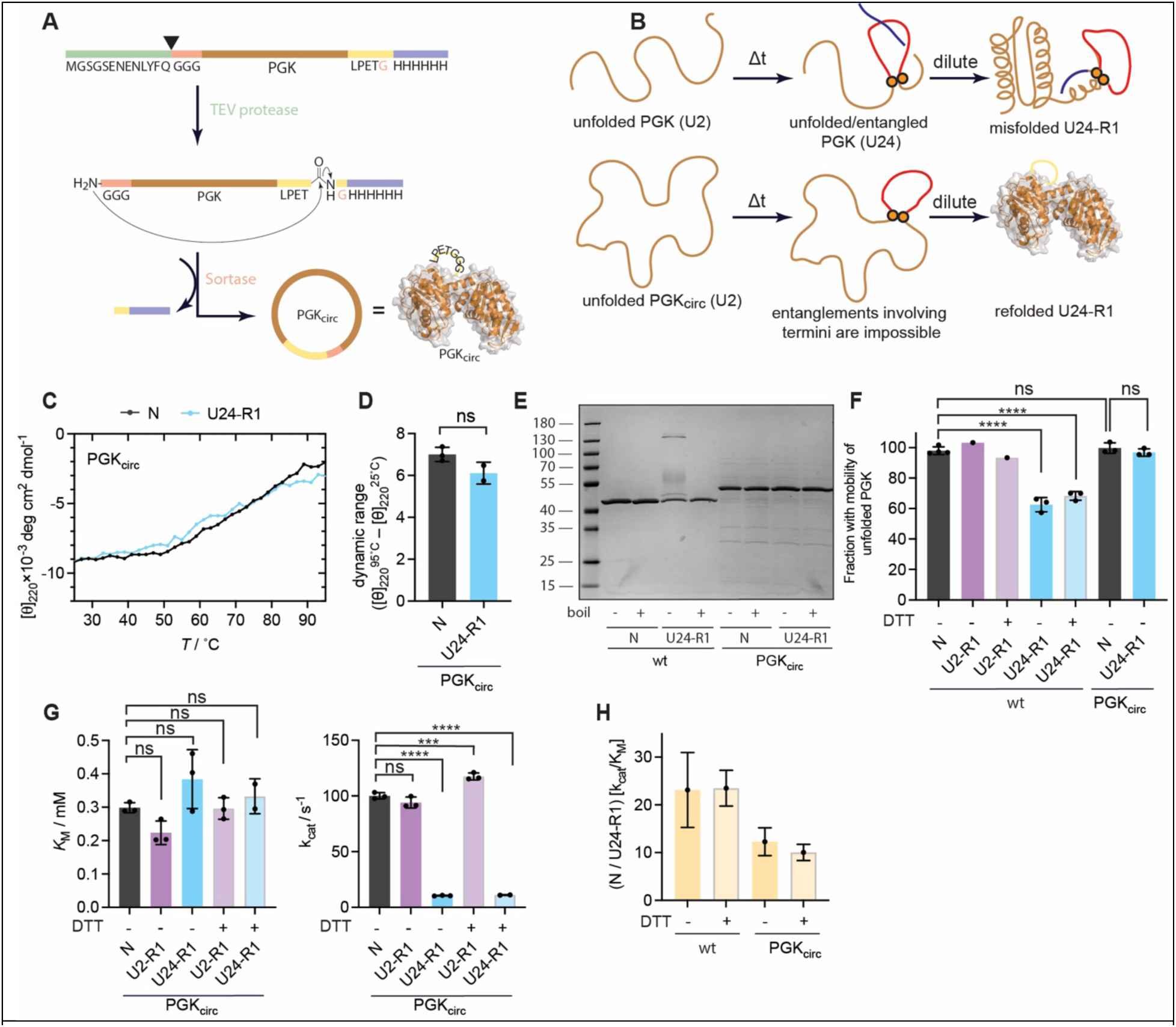
Topological Restriction of PGK Alters its Refolding Outcome. (A) Scheme illustrating the procedure to create a circularized form of PGK (PGK_circ_). (B) Scheme illustrating how circularization could impact refolding by preventing particular conformations from populating in the unfolded ensemble. (C) Thermal melts of PGK_circ_ as monitored by CD signal at 220 nm either on native enzyme or on chemically unfolded-refolded form. (D) Difference in MRE at 95°C endpoint and 25°C starting point for technical replicates (*n* ≥ 2) of the thermal melt. Error bars are std. devs. (E) Native (N) and chemically unfolded-refolded forms of PGK and PGK_circ_ were incubated in 0.6 % SDS, boiled (or not), resolved by Tris-Glycine SDS-PAGE, and visualized with Coomassie stain. (F) Gel image quantification of the fastest-migrating unfolded PGK band in the non-boiled samples relative to the same band in boiled controls for the various forms of PGK and PGK_circ_ with the indicated number of replicates (*n* = 1 to *n* = 3). P-values calculated by one-way ANOVA with Bonferroni’s multiple comparison test. ns, not significant; ****, P < 0.0001. Error bars are std. devs. (G) *K*_M_ and *k*_cat_ derived from fitting replicates (*n* = 3) of Michaelis-Menten kinetics on natively expressed/circularized and chemically unfolded-refolded forms of PGK_circ_. PGK_circ_ was denatured in 6 M GdmCl, with or without 10 mM DTT, for the time indicated. Error bars are std. devs. (H) Comparison of activity loss upon refolding for WT PGK and PGK_circ_.

We next measured the changes in enzymatic activity upon chemically unfolding and refolding PGK_circ_. Compared to wild-type, PGK_circ_’s *k*_cat_ and *K*_M_ were 1.35-fold and 2.82-fold lower, reflecting differences inherent to the circular status, linker sequence, and loss of the His6 epitope tag (Figure 4G). As before, U2-R1 possessed similar catalytic efficiency to the native enzyme (Figure 4G, purple bars) regardless of the presence of reducing agent during denaturation, and U24-R1 possessed a lower catalytic efficiency (Figure 4G, blue bars). However, quantitative comparisons revealed that the extent of activity loss for PGK_circ_ was lower than expected (Figure 4H). Using the catalytic efficiency parameter, *k*_cat_/*K*_M_, wild-type PGK lost (23.1 ± 7.8)-fold of its activity following the U24-R1 treatment whilst PGK_circ_ only lost (12.3 ± 2.9)-fold. Therefore, circularizing PGK prevents some (but not all) of its misfolding pathways, but fully blocks the formation of kinetically stable misfolded states with high resistance to thermal and detergent-induced denaturation.

So far, our data suggest that chemically denatured PGK slowly changes between 2 and 24 h to result in an ensemble that ultimately is less competent to refold. Two possible explanations for this observation are that: (i) U2 is not fully denatured, and hence retains some information of the native state which helps it refold; or (ii) the equilibrated U24 possesses a structural feature that causes the formation of an entanglement when PGK attempts to refold. At a gross level, we found that U2 and U24 both possessed little-to-no secondary structure based on CD (although slight time-dependent differences in the spectra were detectable, Figure 5A) and they also had similar hydrodynamic properties based on their similar sedimentation profiles from AUC (Figure 5B). Small-angle x-ray scattering (Table S2) moreover showed that the unfolded ensemble’s radius of gyration did not change between 2 and 24 h (R_g_ for U2, 79.8 ± 1.4 nm; for U24, 80.6 ± 1.3 nm) and that both the Kratky plots and Flory power-law analysis are consistent with an unfolded, excluded-volume random coil model (Figures S5-S6). It is challenging to obtain detailed structural information about unfolded proteins, however crosslinking mass spectrometry (XL-MS) is suitable for this purpose because nucleophilic residues in PGK that are close to each other spatially can crosslink together upon treatment with a covalent crosslinker even under denaturing conditions (Figure 5C). Hence, we performed a comparative XL-MS experiment in which U2 was crosslinked with light DSBU, U24 was crosslinked with heavy DSBU, and the samples mixed together. Quantifying the relative isotopic intensities for each crosslinked peptide (identified using XiSearch (*59*) and validated with XiFDR (*60*)) thereby reveals residue pairs that get closer together or further apart between U2 and U24. In total, we identified eight unique residue-pairs (Figure 5D). One showed equal abundance at the two timepoints, three were detected only in the PGK that was denatured for 2 h, and two appeared only when PGK was denatured for 24 h (Figure 5D). Among the five crosslinks detected in U2, four were relatively short-distance with respect to sequence, as expected for a self-avoiding random walk polymer because long-distance contacts are higher in entropy (*61*) and would only form preferentially if they are also associated with an attractive interaction. In contrast, the two crosslinks observed at later time points were associated with interactions far from each other along PGK’s primary sequence. Such long-distance interactions could help seed the formation of loops through which other segments could thread during refolding.

**Figure 5.**
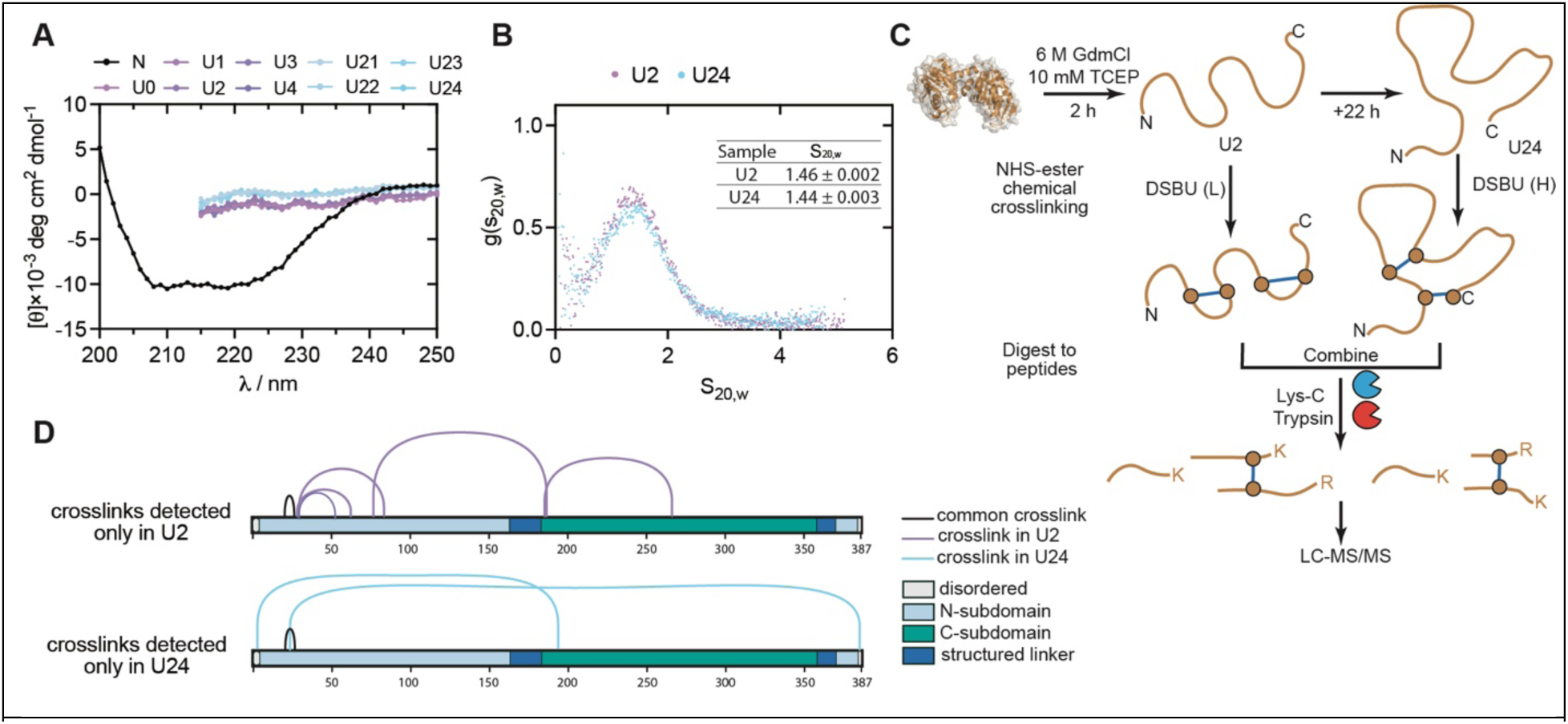
Biophysical and Structural Characterization of PGK’s Unfolded Ensemble. (A) Far-UV circular dichroism (CD) spectra of unfolded PGK, incubated in 6 M GdmCl for the indicated periods of time. Native PGK (black trace) of same material shown as a reference. (B) Representative sedimentation velocity distributions for unfolded PGK, incubated in 8 M urea and 1 mM DTT for the indicated period of time prior to initiating sedimentation. Inset table shows extracted sedimentation velocity and estimated error from fit. (C) Scheme illustrating the crosslinking mass spectrometry (XL-MS) with isotopic labels to assess time-dependent structural evolution of the unfolded ensemble. (D) Connectograms illustrating crosslinks unique to the PGK samples that were unfolded for the indicated periods of time.

We note that the data in Figure 5B-C used reducing agent, because PGK (which contains 3 cysteines) when unfolded in high-concentration denaturant auto-oxidizes with air, forming disulfide linkages. We found this out through the observation that unfolded PGK becomes compacted when it is incubated for 24 h without reducing agent (Figure S7A; AUC of non-reducing) and by Ellman’s assay which shows U24 has fewer reactive thiol groups than U2 (Figure S7B). On the other hand, these disulfide linkages appear to break during refolding and do not alter the outcome of refolding process because: (i) refolded PGK contains the same number of reactive thiols as native PGK (Figure S7B), (ii) all biophysical and enzymatic properties of refolded PGK are insensitive to the presence of reducing agent during unfolding (Figures 3, 4F and S7C).

## Discussion

The model proteins used in classic studies of protein folding were small, soluble, and single-domain, and demonstrated fast reversible refolding kinetics following their denaturation (*62–64*). On the other hand, a number of well-known features associated with primary sequence have been shown to slow down protein folding, such as incorrect disulfide formation (*65–67*) (requiring slow disulfide exchange) and incorrect proline conformation (requiring slow proline isomerization) (*68, 69*). Here we have shown that after being denatured for extended periods of time, PGK cannot spontaneously refold to its native state, and this inefficiency is neither due to disulfide exchange (Figures 3, 4F, and S7C) nor proline isomerization (Figure 6A). Rather, it is due to the fact that PGK forms topologically misfolded states that are very kinetically stable. These states are qualitatively near-native (based on CD spectra and sedimentation velocity). They also cannot be resolved by *E.* coli’s two key chaperones either (GroEL and DnaK) because other work of ours has established PGK as a chaperone-nonrefolding protein as well (*23, 70*). Figure 6B presents a simple model that is consistent with our observations; its key feature is the existence of another section of the energy landscape, separated by high barriers, but accessible from the unfolded ensemble after it has equilibrated. It would be very interesting to know if the state populated by U24-R1 is even more stable thermodynamically than native PGK, akin to alpha lytic protease’s unfolded state (*71*). A long incubation CD experiment showed that even after 72 h, U24-R1’s CD spectrum is unchanged and remains distinct from its native CD spectrum (Figure 6C). Moreover, increasing the refolding time does not result in any appreciable increase in enzymatic activity either (Figure 6D). These experiments show U24-R1 is quite persistent. Whilst these observations can still be consistent with the native state possessing lower Gibbs free energy (given that the barrier between U24-R1 and N could simply be high enough to prevent transitions on this day-timescale), it nevertheless shows that on any timescale relevant to the cell, U24-R1 is irreversibly misfolded.

**Figure 6.**
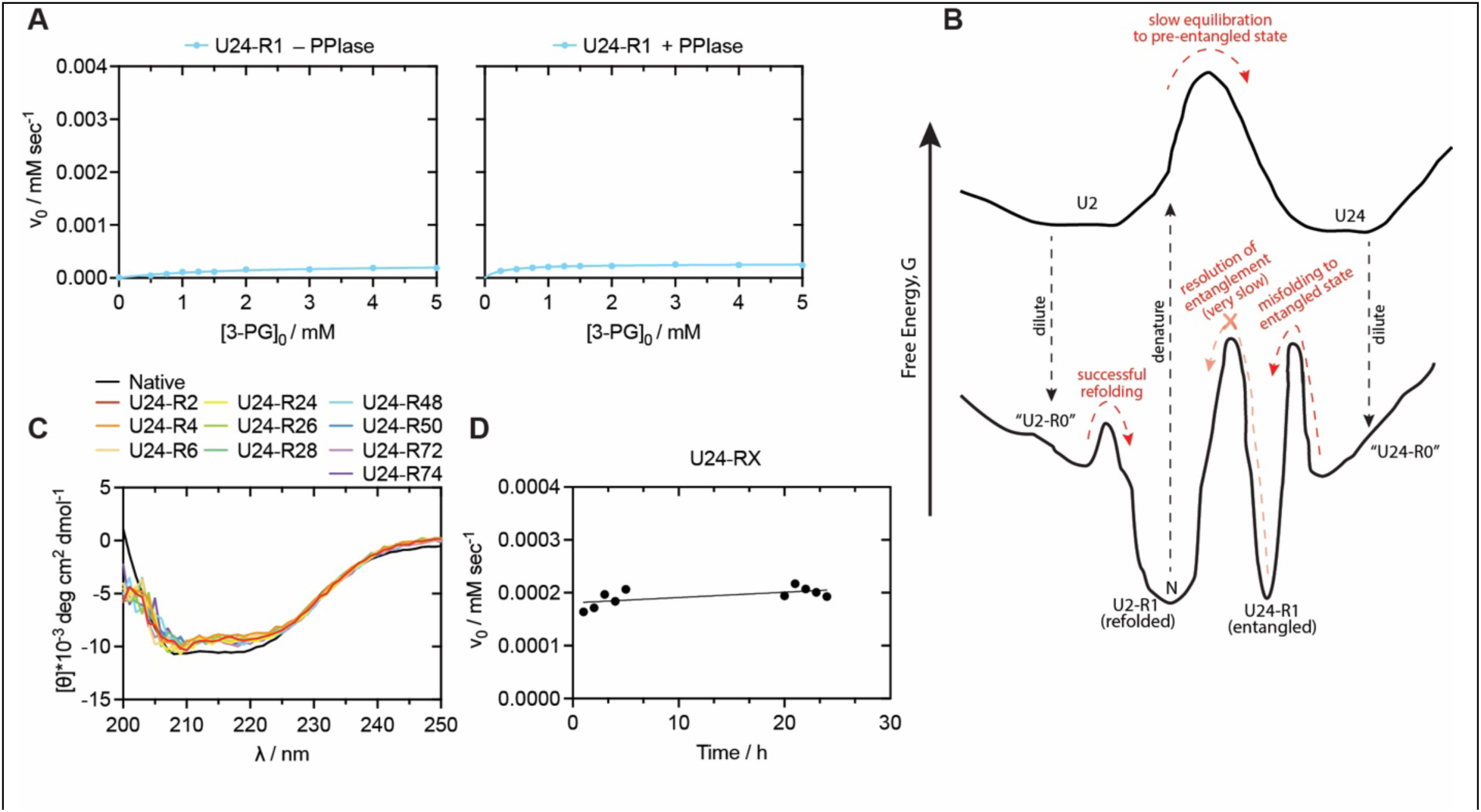
Topologically Misfolded PGK is Persistent. (A) Michaelis-Menten curves showing initial rates as a function of initial substate concentration (3-phosphoglycerate) for (left) native PGK without peptidyl prolyl isomerase (PPIase) and (right) with 1 µM PPIase supplied. (B) Model for the energy landscape of *E. coli* PGK. (C) Far-UV CD spectra of native PGK and chemically unfolded-refolded forms of PGK that were first denatured in 6 M GdmCl, diluted 100-fold, and then incubated for the indicated period of time. (D) Initial enzymatic rates (with 5 mM 3-phosphoglycerate) of chemically unfolded-refolded forms of PGK that were first denatured in 6 M GdmCl, diluted 100-fold, and then incubated for the indicated period of time.

Our experiments raise interesting questions about protein biogenesis and quality control *in vivo*. Protein biophysics tends to emphasize thermodynamic models for protein folding, which characterizes reversible transitions between equilibrium states. However, here we find that a non-equilibrium unfolded ensemble of PGK (U2) is competent to refold, and the equilibrated unfolded ensemble (U24) of PGK is not. Cells conveniently avoid this conundrum because primary protein biogenesis occurs co-translationally, and so, in general nascent proteins never populate full-length equilibrated unfolded ensembles. Therefore, efficient biogenesis of *E. coli* PGK is inherently a non-equilibrium process. We suspect co-translational folding plays an indispensable role for the folding of many proteins (*72–76*); this is probably why cells so often resort to degrading misfolded proteins rather than correcting them.

Although PGK is unlikely to unfold spontaneously on biologically meaningful timescales, AAA+ ATPase chaperones are capable of unfolding folded proteins and can produce expanded ensembles such as those created by denaturant (*77*). As demonstrated here, such processes are risky: they create potential pathways for proteins to misfold into highly stable conformations. In this light, it is perhaps not surprising that most yeast prions are associated with the activity of its versatile AAA+ ATPase Hsp104 (which constitutes both a strength and a liability for yeast proteostasis) (*12*). Moreover, these findings offer a potential reason for why the substrate scope for AAA+ ATPases are so tightly gated and why an all-purpose AAA+ ATPase like Hsp104 has a more limited phylogenetic distribution and disappeared from the metazoan clade (*78*). Whether translocation through AAA+ ATPases could also drive the population of entangled misfolded states like PGK’s U24-R1 would be an interesting subject for future study.

In summary, we have determined the biophysical basis for the nonrefoldability of E. coli PGK. When denatured for extended periods, the protein adopts structural elements which seed the formation of non-native non-covalent lasso entanglements when it attempts to refold. The resulting topologically misfolded state exhibits unusual kinetic stability. It will be important to find out how many other proteins, especially in the human proteome, exhibit this unexpected behaviour.

## Materials and Methods

### PGK expression and purification

The *E. coli pgk* gene with a C-terminal 6×His tag was ordered from Twist as a pre-cloned plasmid, inserted into the pET21(+) vector between the EcoRI and NotI restriction sites. Plasmids were propagated in NEB 10β cells (NEB C3019) as needed. For protein expression, plasmids were transformed into chemically competent NiCo21(DE3) cells (NEB C2527H) by heat shock at 42 °C for 40 seconds in a water bath, recovered in SOC medium at 37 °C for 1 hour, plated on selective LB agar (supplemented with 50 µg/mL ampicillin), and incubated overnight at 37 °C.

Colonies were selected and added to 5 mL starter cultures consisting of Luria-Bertani (LB) broth supplemented with 50 µg/mL ampicillin, and incubated overnight (∼16 h) at 37°C with agitation (220 rpm). The saturated overnight cultures were sub-cultured into 1 L of terrific broth (TB) supplemented with 50 µg/mL ampicillin to a starting OD of 0.05, and were incubate at 37°C with agitation (220 rpm). The cells were induced at OD600 of 0.4 by addition of IPTG to a final concentration of 0.1 mM. After 4 hours of induction, cells were collected by centrifugation (Eppendorf 5910R) at 4000 *g* for 15 min at 4°C in 2× 500 ml centrifuge tubes. The supernatants were discarded, and the resulting cell pellets were flash-frozen in liquid nitrogen and then stored at -20°C.

For protein purification, cell pellets were resuspended in 25 mL lysis buffer (20 mM HEPES-NaOH pH 7.4, 100 mM NaCl, 2 mM MgCl_2_, 20 mM imidazole-HCl pH 8.0). The protease inhibitor, phenylmethylsulfonyl fluoride (PMSF), was added to a final concentration of 1 mM, and Deoxyribonuclease I (DNase I) was added to a final concentration of 0.1 mg/ml. The mixture was subsequently homogenized using a QSonica sonicator (amplitude = 55%, cycling between 8 s on, 8 s off, for 15 min total on-time) on an ice-water slurry. The lysate was clarified by centrifugation at 16,000 *g* for 15 min at 4°C and then purified by Ni-NTA chromatography. First, clarified lysate was combined with 2 mL of Ni-NTA resin (preequilibrated into lysis buffer) in a column with its ends sealed, and allowed to incubate for 1 h at 4°C with gentle rocking. The column was then set vertically and unsealed to allow lysate to flow through the Ni-NTA bed. The beads were subsequently washed with 4 x 10 mL wash buffer (20 mM HEPES-NaOH pH 7.4, 100 mM NaCl, 2 mM MgCl_2_, 40 mM imidazole-HCl pH 8.0), and then eluted with 3 mL elution buffer (20 mM HEPES-NaOH pH 7.4, 100 mM NaCl, 2 mM MgCl_2_, 300 mM imidazole-HCl pH 8.0) which was passed through the Ni-NTA bed twice.

Eluted proteins were analyzed by 4-20% Tris-Glycine SDS-PAGE and intact mass spectrometry to assess purity and confirm molecular weight. As a single band at the expected molecular weight was observed by Coomassie staining and confirmed by native mass spectrometry, no further purification was performed. The eluted proteins were dialyzed into native buffer (20 mM HEPES-NaOH pH 7.4, 100 mM NaCl, 2 mM MgCl_2_) overnight using G2 dialysis cassettes, 10k MWCO (Fisher Scientific).

Protein concentration was determined by BCA assay (Thermo Fisher) following the manufacturer’s protocol (typical concentration would be ∼14 mg/ml). Two protein stocks were prepared each to a final concentration of 10 mg/mL. Stock 1 – which is used to generate “native samples” – was prepared by adding the appropriate volume split between 100% glycerol and native buffer to get to final concentrations of 10% (v/v) glycerol and 10 mg/ml protein. Stock 2 – which is used to generate “refolded samples” – was prepared by adding the same volume of native buffer. Both stocks were aliquoted out into 50 µL aliquots in microfuge tubes, flash frozen in liquid nitrogen, and stored at -80°C until future use.

### Native and refolded sample preparation

To prepare native samples for downstream experiments, Stock 1 was diluted 100-fold using native dilution buffer (20 mM HEPES-NaOH pH 7.4, 100 mM NaCl, 2 mM MgCl₂, and 0.06 M guanidinium chloride (GdmCl)), so that typical final conditions for native samples are 20 mM HEPES-NaOH pH 7.4, 100 mM NaCl, 2 mm MgCl_2_, 0.0594 M GdmCl, 0.1% (v/v) glycerol, 0.1 mg/ml PGK. Sample volumes were adjusted as needed for each experiment.

Refolded samples were prepared by taking a defined amount of protein from Stock 2 (depending on the experimental setup) and drying it completely using a Vacufuge centrifugal concentrator (Eppendorf). To initiate unfolding, an equal volume of freshly prepared unbuffered 6 M GdmCl solution was added to the dried protein+salt pellet. The mixture was resuspended by pipetting up and down and incubated at room temperature for a defined period of time. Refolding was initiated by diluting the unfolded protein 100-fold with refolding dilution buffer (20 mM HEPES-NaOH pH 7.4, 100 mM NaCl, 2 mM MgCl₂, and 0.1% (v/v) glycerol), so that typical final conditions for refolded samples are 20 mM HEPES-NaOH pH 7.4, 100 mM NaCl, 2 mM MgCl_2_, 0.06 M GdmCl, 0.099% (v/v) glycerol, 0.1 mg/ml PGK. PGK was incubated for the indicated periods of time at room temperature (typically 1 h) before commencing downstream experiments. The resulting samples are referred to as U2-R1 or U24-R1, depending on the duration of the unfolding incubation (2 h or 24 h, respectively).

For other experiments, unfolding was conducted in (a) 8 M urea, (b) 6 M GdmCl, 10 mM DTT. (Case a): To initiate unfolding, an equal volume of freshly prepared unbuffered 8 M urea solution was added to the dried protein+salt pellet. The mixture was resuspended by pipetting up and down and incubated at room temperature for a defined period of time. Refolding was initiated by diluting the unfolded protein 100-fold with refolding dilution buffer (20 mM HEPES-NaOH pH 7.4, 100 mM NaCl, 2 mM MgCl₂, and 0.1% (v/v) glycerol), so that typical final conditions for refolded samples are 20 mM HEPES-NaOH pH 7.4, 100 mM NaCl, 2 mM MgCl_2_, 0.08 M urea, 0.099% (v/v) glycerol, 0.1 mg/ml PGK. (Case b): To initiate unfolding, an equal volume of freshly prepared unbuffered 6 M GdmCl, 10 mM DTT solution was added to the dried protein+salt pellet. The mixture was resuspended by pipetting up and down and incubated at room temperature for a defined period of time. Refolding was initiated by diluting the unfolded protein 100-fold with refolding dilution buffer (20 mM HEPES-NaOH pH 7.4, 100 mM NaCl, 2 mM MgCl₂, and 0.1% (v/v) glycerol), so that typical final conditions for refolded samples are 20 mM HEPES-NaOH pH 7.4, 100 mM NaCl, 2 mM MgCl_2_, 0.06 M GdmCl, 0.1 mM DTT, 0.099% (v/v) glycerol, 0.1 mg/ml PGK.

### Circular dichroism experiments

Circular dichroism (CD) experiments were carried out on a Jasco J-810 spectropolarimeter using 250 µL of 0.1 mg/mL protein samples typically prepared in 20 mM HEPES-NaOH pH 7.4, 100 mM NaCl, 2 mM MgCl₂, 0.06 M guanidinium chloride (GdmCl), and 0.1% (v/v) glycerol.

Baseline CD spectra were collected at 25 °C using a 1 mm pathlength quartz cuvette over a wavelength range of 200–250 nm. Spectra were corrected by subtracting the signal from a matched buffer blank. Thermal denaturation was monitored by measuring ellipticity at 220 nm from 25 °C to 90 °C in 1 °C increments, using the same buffer conditions. After thermal ramping, samples were cooled back to 25 °C, and a second CD spectrum was recorded to assess structural reversibility.

### SDS PAGE detergent sensitivity assay

Protein samples were analyzed on 4-20% Tris-Glycine SDS-PAGE gels (Thermo Fisher). For a typical sample, 20 µL of PGK (either native or chemically unfolded-refolded, 0.1 mg/ml) was combined with 1.33 µL of concentrated loading buffer (0.35 M Tris-HCl (pH 6.8), 30% (v/v) glycerol, 10% SDS, 0.6 M DTT, and 1.5 mM bromophenol blue). After combining, final SDS concentration is 0.62%. Next, samples were either boiled at 90 °C for 10 minutes or allowed to sit at room temperature, and then 20 µL were loaded into wells of the polyacrylamide gel. Electrophoresis was performed at 180 V for 1 hour in Tris-Glycine running buffer containing 25 mM Tris, 192 mM glycine, 0.33% SDS. Gels were stained with Imperial Coomassie stain (Thermo Fisher) for 20 minutes and destained in deionized water for 1 hour. Gel images were captured using a Bio-Rad ChemiDoc imaging system and analyzed in ImageJ. Image analysis was performed by measuring the integrated density associated with equally-sized boxes centered on gel bands and taking the appropriate ratios, without any image manipulation or background subtraction.

### Standard SDS-PAGE (not for detergent sensitivity assays)

Protein samples were analyzed on 4-20% Tris-Glycine SDS-PAGE gels (Thermo Fisher). For a typical sample, 20 µL of sample (e.g., fractions from purification steps) was combined with 4 µL of concentrated loading buffer (0.35 M Tris-HCl (pH 6.8), 30% (v/v) glycerol, 10% SDS, 0.6 M DTT, and 1.5 mM bromophenol blue). After combining, final SDS concentration is 1.66%. Samples were boiled at 90 °C for 10 minutes, and then 20 µL were loaded into wells of the polyacrylamide gel. Electrophoresis was performed at 180 V for 1 hour in Tris-Glycine running buffer containing 25 mM Tris, 192 mM glycine, 1.0% SDS. Gels were stained with Imperial Coomassie stain (Thermo Fisher) for 20 minutes and destained in deionized water for 1 hour. Gel images were captured using a Bio-Rad ChemiDoc imaging system.

### Enzyme Kinetics Assay

The enzymatic activity of native PGK, U2-R1, and U24-R1 was measured at room temperature using a GAPDH-coupled spectrophotometric assay as previously described(*79*). Reactions were conducted in a final volume of 200 µL containing 20 mM HEPES-NaOH pH 7.4, 100 mM NaCl, 2 mM MgCl₂, 0.2 mM NADH, 0.5 mM EDTA, 5 mM MgATP, 0.4 μM GAPDH (Millipore Sigma, G2267), and varying concentrations (0–5 mM) of 3-phosphoglycerate (3-PG). Reactions were initiated by the addition of native PGK or a chemically-refolded PGK (previously unfolded and refolded for various intervals of time) to a final concentration of 0.0325 µM. Kinetic parameters were determined at fixed ATP and NADH concentrations, whilst 3-PG concentration was varied to obtain Michaelis-Menten curves. All measurements were acquired on a Tecan Spark microplate reader by measuring absorbance at 340 nm every 20 s for 15 min. Initial velocities were derived from the slope of the linear portion of the time series, and then dividing by the slope of an NADH calibration curve (to convert from A_340_/min to [NADH]/min). Unimolecular rate constants were then derived by dividing maximal rate by enzyme concentration.

### Crosslinking Experiments on Unfolded PGK

Unfolded PGK samples were prepared by taking 2 µL of Stock 2, drying it completely using a Vacufuge centrifugal concentrator, and adding 2 µL of 6 M GdmCl, 10 mM TCEP to the dried pellets+salt. PGK was incubated for either 2 h or 24 h, and then diluted 100-fold with 198 µL of unfolding buffer (20 mM HEPES-NaOH pH 7.4, 100 mM NaCl, 2 mM MgCl₂, 6 M GdmCl). To the 200 µL, 2 µL of crosslinker was added, either DSBU-d0 (100 mM stock in DMSO; ThermoFisher A35459) to U2, or DSBU-d12 (a deuterated analog, 100 mM stock in DMSO, synthesized in-house (*37*)) to U24. Crosslinking reactions were incubated at room temperature for 1 h on a Roto-Mini rotator (10 rpm). Crosslinking was quenched by adding 1 M Tris-HCl, pH 7.5 to a final concentration of 20 mM, followed by an additional 30-min incubation at room temperature with rotation (10 rpm). Crosslinking was performed in three replicates for both native and refolded PGK, pairs of U2 and U24 were mixed, resulting in three samples.

### Mass Spectrometry Sample Preparation

Crosslinking samples were reduced by incubation with 10 mM DTT at 37 °C for 30 min with agitation on a thermomixer. Free thiols were subsequently alkylated by incubation with 40 mM iodoacetamide at room temperature for 45 min in the dark.

Lys-C digestion (NEB P8109S) was initiated by adding 4 µL of 0.1 mg/mL Lys-C (1:100 w/w enzyme-to-substrate ratio), followed by incubation at 37 °C and 700 rpm for 1.5 h. After Lys-C digestion, reactions were diluted four-fold with 100 mM ammonium bicarbonate, pH 8.0.

Trypsin digestion (NEB P8101S) was performed by adding 8 µL of 0.1 mg/mL trypsin (1:50 w/w enzyme-to-substrate ratio), followed by overnight incubation at 25 °C and 700 rpm.

Peptide mixtures were acidified with trifluoroacetic acid (TFA, Acros) to a final concentration of 1% (v/v) and desalted using Sep-Pak C18 1cc Vac cartridges (Waters). Cartridges were preconditioned with 2 mL of buffer B (80% acetonitrile, 0.5% TFA in LC-MS grade water) and equilibrated with 4 mL of buffer A (0.5% TFA in LC-MS grade water). Acidified peptides were loaded under vacuum, washed with 4 mL of buffer A, and eluted with 1 mL of buffer B. Eluates were dried using a Vacufuge concentrator (Eppendorf) and stored at - 80 °C until LC-MS analysis.

### LC-MS/MS Acquisition

LC-MS/MS analysis was performed using a Thermo UltiMate 3000 UHPLC system coupled to a Q Exactive HF-X Orbitrap mass spectrometer (Thermo Fisher Scientific). Peptides were separated on an Acclaim PepMap RSLC C18 column (75 µm × 25 cm, 2 µm, 100 Å) at 300 nL/min and 40 °C.

Approximately 1 µg of peptide digest was loaded onto a trap column and concentrated for 10 min at 2% buffer B (A: 0.1% formic acid in water; B: 0.1% formic acid in acetonitrile). For crosslinked samples, a 75-min gradient was applied: 2% B over 10 min, 2-35% over 45 min, 35-40% over 5min, and 40–90% over 5 min, followed by a 5 min column wash.

Data were acquired in positive ion mode using a data-dependent top-20 method. Full MS scans (m/z 300–1800) were acquired at 120,000 resolution with a 3e6 AGC target and 100 ms injection time. MS/MS scans were collected at 60,000 resolution, AGC target of 2e5, 250 ms injection time, and 2 m/z isolation window. Precursors showing the characteristic 12.075 Da d0/d12 mass shift were prioritized for fragmentation. Stepped HCD was applied (NCEs 22, 25, 28). Ions with charges of 1, 2, or > 8 were excluded; dynamic exclusion was set to 60 s.

### Data analysis for crosslinking experiment

The Proteome Discoverer Software Suite (PD, v2.4, Thermo Fisher) in conjunction with the Minora feature detector algorithm were employed to conduct spectral search and label free quantification (LFQ) of the peptides identified.

The LFQ analyses included three replicate crosslinked raw files. We used the MSFragger node for peptide identification set to perform a standard-tryptic search and allowing up to 2 missed cleavages. The precursor mass tolerance was set to 10 ppm at the MS1 level and a fragment ion tolerance of 0.02 Da was set for the MS/MS level. We allowed oxidation of methionine and acetylation of the N-terminus as dynamic modifications, while carbamidomethylation on cysteines was set as a static modification. The search was conducted using a FASTA file that included just the sequence for E. coli PGK (P0A799).

Although we do not use PD to identify crosslinked peptides, PD produces an output called a consensus feature file which consists of all detected ion intensities for each feature (annotated by its precursor m/z and retention time) regardless of whether they are assigned to a peptide spectrum match (PSM) or not. We used these consensus feature outputs to obtain ion intensities for crosslinked peptides that were confidently assigned to spectra using XiSearch (*59*).

To identify crosslinks in XiSearch, the raw data were first converted to .mgf files and recalibrated with MSConvert. The standard “small-scale” setting of searching for DSBU crosslinks from the software was used. Crosslink specificity sites were defined as lysine and protein N-termini as site1, and lysines, protein N-termini, serines, threonines, and tyrosines as site2. In the settings file, methionine oxidation was treated as a variable modification, while cysteine carbamidomethylation was enforced as a fixed modification. XiSearch was configured to search only b- and y-ions, and the digestion enzyme was set to trypsin. Both water loss and ammonia loss were allowed. We permit up to three missed cleavages and two modifications per peptide. Mass deviations of up to 5 ppm (MS1 level) and 10 ppm (MS/MS level) were allowed.

XiSearch was run by supplying the settings file (described above), .mgf files, and the FASTA file consisting of just the sequence for E. coli PGK (P0A799) as inputs. We next ran a separate XiSearch that used the same settings as above but in which the crosslinker was adjusted from standard DSBU to DSBU-d12. In the settings file, DSBU-d12 was implemented under advanced config by supplying the crosslinker’s name and mass (208.15979231 Da; mass difference of 12.075 Da from DSBU-d0). After the data was processed by XiSearch, we provided separate output data files (one from search against DSBU-d0, one from search against DSBU-d12) to XiFDR (71), along with the settings file (from XiSearch) and the FASTA file. XiFDR was used with its default settings with one modification, a prefilter for doublet counts (CCfragmentdoubletcount) to be greater than 0. Then, we concatenated the XiFDR output files for DSBU-d0 and DSBU-d12 searches.

To process the data (namely, the assignments from XiSearch/XiFDR and the quantifications from PD consensus feature files), we used custom Python scripts, described in the following and available upon request.

We start with a list of confident crosslinked peptide-spectrum matches (XSMs) identified from XiSearch/XiFDR. Each of these crosslinked peptides are annotated with which injection they came from, scan ID, retention time, and a precursor m/z. Using this information, we identify which feature in the PD consensus feature file maps to the crosslinked peptide precursor (based on m/z and retention time). This provides three ion intensities for the particular crosslinked peptide in the three replicates. Missing values are filled with an imputed ion count of 10_3_, representing an estimated detection limit.

If the crosslinked peptide was crosslinked with DSBU-d0, the script attempts to identify a consensus feature that would correspond to the same species except crosslinked with DSBU-d12. To do this, it looks for a feature with a matching retention time (within a 3 min tolerance) and a precursor m/z that is greater by 12.075 divided by the charge state (within a 10 ppm tolerance). If no matching consensus feature is found, we treat the heavy crosslinked peptide as absent and fill its values with an imputed ion count of 10_3_. However, if a matching consensus feature is found, the associated ion intensities are extracted and grouped together with the ion intensities for the DSBU-d0 replicates. If the crosslinked peptide was crosslinked with DSBU-d12 the script performs the same function except it looks for a consensus feature that would correspond to the same species crosslinked with DSBU-d0. This requires matching retention time (±3 min) and a precursor m/z that is less by 12.075 divided by the charge state (±10 ppm).

Next, we calculated the ratio associated with each consensus feature by dividing the total native extracted ion intensities by the total refolded extracted ion intensities. Each unique crosslink (as represented as a residue-pair) could be observed through multiple consensus features: this could arise through different peptides getting crosslinked together at the same two residues, or the same crosslinked peptide being observed in different charge states, methionine oxidation states. To report a single quantification for each unique crosslink, we group together the data from all the consensus feature that map to the same unique crosslink.

Specifically, if the ratios associated with the consensus features disagree in sign (e.g., if a crosslinked peptide is more abundant in U2 in a 2+ charge state, but in a 3+ charge state, it’s more abundant in U24), the crosslink is filtered out. A “majority rules” heuristic is used so that if two consensus features agree in sign and one does not, the disagreeing feature is discarded and the remaining two are retained.

### Construction of the PGK-Circ Plasmid

The sequence GSGSENENLYFQGGG was introduced between the initial methionine and the second amino acid of PGK by QuickChange inverse PCR on the PGK plasmid cloned in the pET-21(+) vector. PCR products were treated with 0.5 µL DpnI for 30 min at 37 °C and purified using the Zymo DNA Clean & Concentrator Kit (Zymo Research) following the manufacturer’s protocol. DNA size was verified by gel electrophoresis, and concentration was measured using a NanoDrop One UV-Vis spectrophotometer (Thermo Scientific).

Purified PCR product was transformed into NEB 10-beta chemically competent *E. coli* by heat shock and plated on LB agar supplemented with 100 µg/mL ampicillin. Several selected colonies were grown overnight in 5 mL LB with 100 µg/mL ampicillin at 37 °C and 220 rpm. Plasmids were extracted using a mini-prep kit (Zymo) and verified by Sanger sequencing.

For C-terminal modification, the verified altered N-terminal construct was used as a template to introduce the sequence GGGSGGGSLPETG before the His-tag using the same mutagenesis and cloning procedure. The final construct encoded MGSGSENENLYFQGGG-PGK-GGGSGGGSLPETG-HHHHHH, incorporating short peptide adducts at both termini.

### PGK Circ expression and purification

PGK_circ_ was expressed following the same protocol as wild-type PGK described above. Purification steps were identical up to the elution stage. Elution was performed using a stepwise imidazole gradient (70 mM, 150 mM, and 300 mM). The final eluate was further purified by passing through a chitin affinity column, and the flow-through was collected. Protein purity was assessed by 4–20% SDS-PAGE. The purified protein was buffer-exchanged into 20 mM HEPES-NaOH pH 7.4, 100 mM NaCl, 2mM MgCl_2_ using 10k MWCO centrifugal concentrators (Amicon, Millipore). Protein concentration was determined using the BCA assay (Thermo Fisher Scientific) according to the manufacturer’s instructions. The protein was concentrated to 1 mg/mL, supplemented with 20% glycerol, re-quantified by BCA assay, flash-frozen in liquid nitrogen, and stored at -80 °C until use.

### PGK_circ_ Circularization and Intermolecular Transpeptidation

The pre-PGK_circ_ construct, MGSGSENENLYFQ-GGG-PGK-LPETG-HHHHHH, was circularized in a two-step reaction. First, TEV protease (NEB, P8112S) was added to a final concentration of 0.74 μM (1:50 w/w enzyme-to-substrate ratio) and incubated for 1 hour at room temperature to cleave the N-terminal tag (MGSGSENENLYFQ), leaving behind an exposed N-terminal GGG motif.

Next, CaCl_2_ was spiked into the solution to a final concentration of 1 mM, and Sortase A (BPS Bioscience) was added to a final concentration of 0.4 μM. The reaction was incubated at 37 °C for 24 h. Following the reaction, the mixture was subjected to inverse Ni-NTA resin chromatography, by incubating with a bed of Ni-NTA agarose and collecting the flow-through. Circularization was confirmed by SDS-PAGE and intact mass spectrometry.

Purified PGK_circ_ was buffer exchanged into 20 mM HEPES-NaOH (pH 7.4), 100 mM NaCl, and 2 mM MgCl₂, and concentrated to 10 mg/mL. For future use, the protein was either stored in buffer alone (to generate “Stock 2”) or supplemented with 10% glycerol (to generate “Stock 1”) following the same protocol as for wild-type PGK. PGK_circ_ was flash-frozen in liquid nitrogen and stored at -80 °C until use.

### Quantification of Free Cysteine Residues Using Ellman’s Assay

Free cysteine content in PGK samples (native, U2, U24, U2-R1, and U24-R1) was quantified using the standard Ellman’s assay. To prepare unfolded PGK samples, 2 µL of Stock 2 was dried completely using a Vacufuge centrifugal concentrator. The dried pellet (including salts) was then resuspended in 2 µL of 6 M guanidinium chloride (GdmCl), incubated at room temperature for either 2 h (U2) or 24 h (U24), and then diluted 100-fold with 198 µL of unfolding buffer (20 mM HEPES-NaOH, pH 7.4, 100 mM NaCl, 2 mM MgCl₂, 6 M GdmCl). After unfolding, 60 µL of each sample was transferred to a 96-well plate and mixed with 140 µL of assay buffer containing 5,5′-dithiobis-(2-nitrobenzoic acid) (DTNB) at a concentration that yields a final DTNB concentration of 0.18 mM in the 200 µL reaction. The mixture was incubated at room temperature for 15 min, and absorbance at 412 nm was recorded on a Tecan Spark microplate reader to monitor the formation of 2-nitro-5-thiobenzoic acid (TNB). Refolded PGK samples (U2-R1 and U24-R1) were prepared by diluting unfolded U2 and U24 samples 100-fold into native buffer (20 mM HEPES-NaOH, pH 7.4, 100 mM NaCl, 2 mM MgCl₂, 0.1% (v/v) glycerol) and incubating at room temperature for 1 h. Ellman’s assay was then carried out as described above, using 60 µL of refolded sample and 140 µL of DTNB-containing assay buffer. Native PGK was assayed in parallel under the same conditions at 0.1 mg/mL. A standard curve was generated using a fixed concentration of DTNB and a series of dithiothreitol (DTT) dilutions corresponding to 1, 2.5, 5, 10, 25, and 50 µM cysteine equivalents, in a total volume of 200µL. Absorbance at 412 nm was used to establish a linear correlation between TNB formation and free thiol concentration.

### Sedimentation Velocity Analytical ultracentrifugation Sample Preparation

Analytical ultracentrifugation (AUC) experiments were conducted using a Beckman Coulter XL-A analytical ultracentrifuge. Unfolded PGK samples were prepared by taking 6 µL of Stock 2, drying it completely using a Vacufuge centrifugal concentrator, and resuspending the dried pellet (including salts) in 6 µL of 8 M urea with or without 1 mM DTT. Samples were incubated for either 2 h or 24 h, then diluted 100-fold with 594 µL of unfolding buffer (20 mM HEPES-NaOH, pH 7.4, 100 mM NaCl, 2 mM MgCl₂, 8 M urea, with or without 1 mM DTT). To assess the structural properties of chemically unfolded-refolded samples (U2-R1 and U24-R1), the dried pellet was resuspended in 6 µL of GdmCl, incubated for either 2 h or 24 h, then diluted 100-fold with 594 µL of refolding buffer, as described above.

### Sedimentation Velocity Analysis by Analytical Ultracentrifugation

The Sedimentation constant was determined for PGK under a variety of conditions using sedimentation velocity analytical ultracentrifugation (SV-AUC). All SV-AUC experiments were performed using a Beckman XL-A ultracentrifuge (Beckman Coulter) and samples were immediately loaded into cells with 1.2 mm double-sector epoxy centerpieces and sapphire windows. Each sample was spun at 25°C using a 4-hole, An-Ti60 rotor and speed of 50,000 rpm for up to 400 scans. Radial scans were acquired with 0.003 cm radial steps in continuous mode no delay between scans at a wavelength of 235 nm.

Weight-average sedimentation coefficient distributions, g(s*sw) were created using dc/dt+ (80) and were corrected using the appropriate densities (ρ), viscosities (η), and partial specific volumes (n) for each buffer and protein content and was calculated using SEDNTERP version 3.0.4 (Table S1) (81).

### Chemical Denaturation Experiments

Chemical denaturation of PGK was monitored by circular dichroism (CD) at 220 nm at 25 °C across a range of guanidinium chloride (GdmCl) concentrations from 0 to 2.5 M (35 total concentrations). Samples were incubated for 48 hours prior to measurement to ensure equilibration at each denaturant concentration. The global thermodynamic stability was determined by fitting the data to a two-state unfolding model (*82*).

### Time-dependent SAXS data collection

Small angle X-ray solution scattering (SAXS) data were collected at Sector 7A1 of the Cornell High Energy Synchrotron Source using an 11.21 keV, 250 µm beam with flux of 3.58 × 10_12_ photons/s. Dried protein samples were dissolved in buffer containing 8 M urea and spun at 14,000 rpm for 5 min prior to data collection. For each measurement, a 30 µl volume of sample was oscillated in a 1.5 OD quartz capillary while ten 1 s exposures were collected twice in succession. Scattering images were collected at 23 °C and captured on an EIGER 4M detector (Dectris, Switzerland), normalized using integrated beamstop diode counts. All exposures were averaged using the CorMap algorithm (*83*) implemented in the RAW data processing software (*84*) to detect any progressive radiation damage. Complete SAXS acquisition parameters are listed in Table S2.

Due the time resolved nature of this experiment and the presence of concentrated urea, we obtain matching buffer by special procedure: a 500 µl 10k MWCO Amicon centrifugal concentrator was pre-rinsed with 200 µl of 0.1 M NaOH followed by two rinses of 200 µl of buffer (8 M urea), then spun to dryness to remove any preservatives and residual liquid. Half of every sample volume was set aside from data collection (unexposed) and spun through the pre-rinsed concentrator so that the filtrate would serve as matched buffer. Each sample was prepared in duplicate. Samples prepared the day before data collection served as the “24 h” time points (actual elapsed time to completion: 26 h 05 min). Samples prepared same-day, “2 h”, were completed after 2 h 29 min.

## Acknowledgments

The authors acknowledge Drs. Yang Jiang and Edward O’Brien for thoughtful comments and feedback. The authors acknowledge Dr. Eugene Shakhnovich for proposing the idea of circularizing PGK. The authors thank Phil Mortimer for supporting the mass spectrometry core facility at Johns Hopkins Chemistry, and Katherine Tripp for training and support with circular dichroism experiments. This work is based on research conducted at the Center for High-Energy X-ray Sciences (CHEXS).

## Funding

National Science Foundation MCB-2045844 (SDF)

National Institutes of Health DP2-GM140926 (SDF)

Camille Dreyfus Teacher-Scholar Award (SDF)

Sloan Fellowship (SDF)

Ada Sinz Fellowship (YX)

National Science Foundation MCB-2427621 (KGF)

National Institutes of Health R35-GM148199 (KGF)

National Science Foundation DMR-2342336 (REG)

National Institutes of Health 1-P30-GM124166 (REG)

## Author contributions

Conceptualization: YX, SDF

Methodology: YX

Investigation: YX, BA, REG, PS

Formal Analysis: YX, BA, REG

Visualization: YX, SDF

Supervision: KGF, SDF

Writing—original draft: YX, SDF

Writing—review & editing: YX, BA, KGF, SDF

Funding Acquisition: REG, KGF, SDF

Project Administration: SDF

## Competing interests

The authors declare no competing interests.

## Data and materials availability

All other data needed to evaluate the conclusions in this paper are present in the paper and/or the Supplementary Materials

## Supplementary Materials

**This PDF file includes:**

**Fig. S1.**
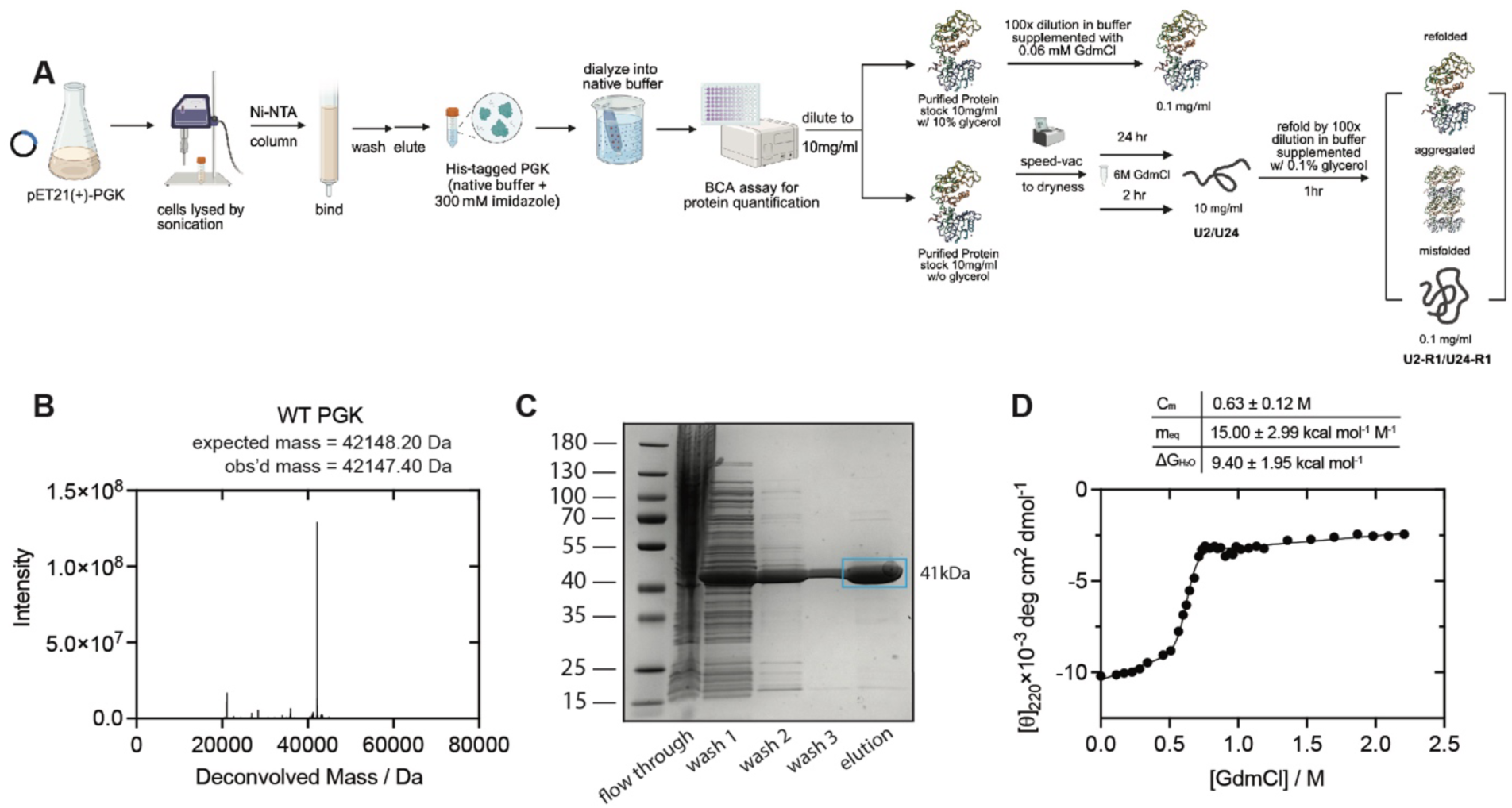
Purification and Characterization of Wild-Type PGK. (A) Scheme illustrating the purification and storage of wild-type PGK in two stocks, later used for native, U2-R1, and U24-R1 samples. (B) Max-entropy deconvolved intact mass spectrum confirming the molecular weight of the purified PGK. (C) SDS-PAGE stained with Coomassie showing pure PGK in the elution fraction. (D) GdmCl-induced unfolding curve of wild-type E. coli PGK. Continuous line represents a two-state model fit. The global free energy of unfolding is 9.40 kcal/mol in 20 mM HEPES pH 7.4, 100 mM NaCl, 2 mM MgCl_2_ at 25°C.

**Fig. S2.**
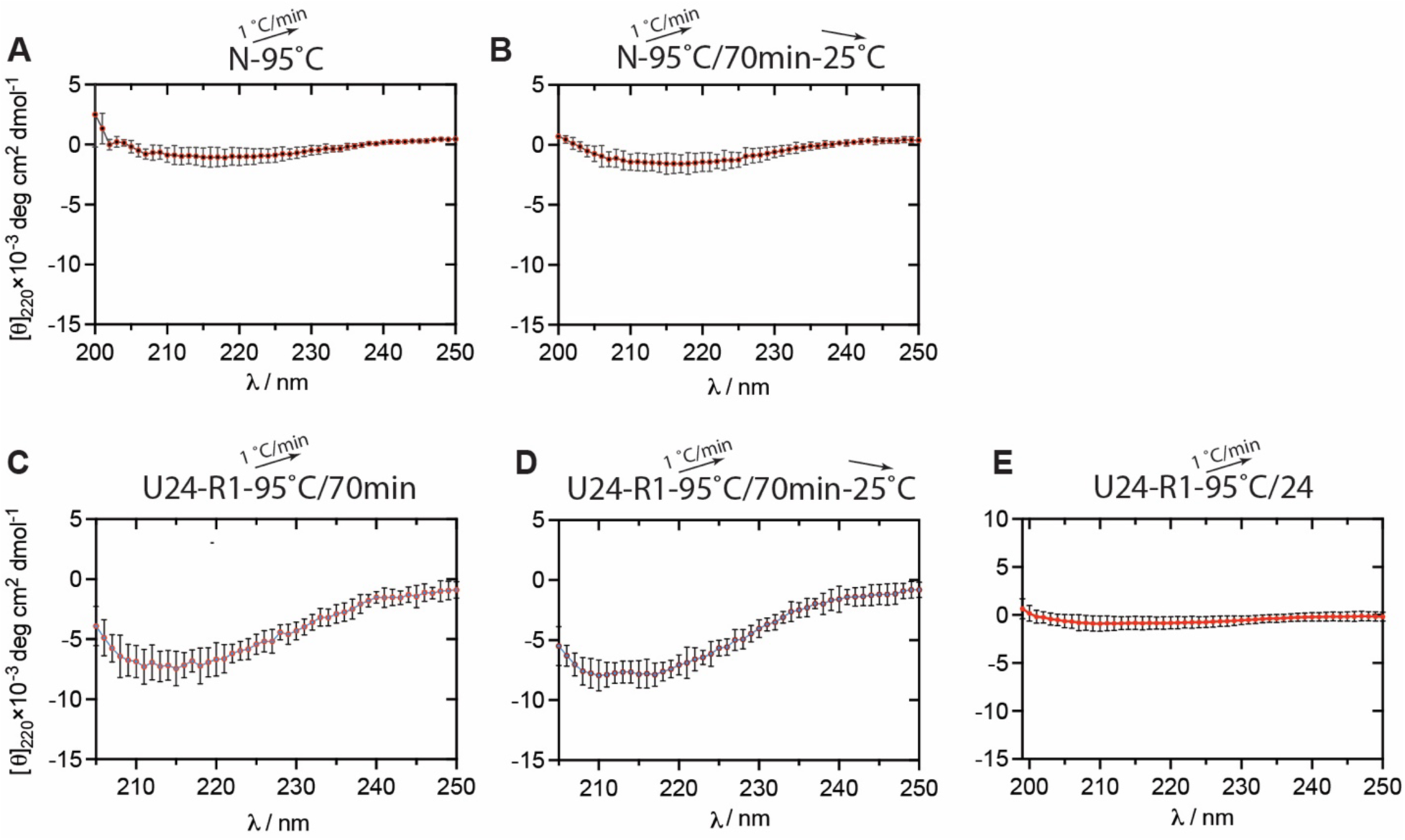
Thermal Denaturation of Native and Refolded PGK. For all panels, n = 3; dots represent means; error bars represent std. devs. All spectra were acquired on 0.1 mg/ml PGK in 20 mM HEPES-NaOH pH 7.4, 100 mM NaCl, 2 mm MgCl_2_, 0.06 M GdmCl, 0.1 % (v/v) glycerol, 0.1 mg/ml PGK. (A) Ending CD spectrum of native PGK after ramping upto 95°C at 1 °C/min. (B) Ending CD spectrum of native PGK after ramping upto 95°C at 1 °C/min and then cooling to room temperature on benchtop. (C) Ending CD spectrum of PGK that was first denatured in 6 M GdmCl for 24 h, then diluted 100-fold and incubated for 1 h to refold, and then ramping upto 95°C at 1 °C/min. (D) Ending CD spectrum of PGK that was first denatured in 6 M GdmCl for 24 h, then diluted 100-fold and incubated for 1 h to refold, then ramping upto 95°C at 1 °C/min and then cooling to room temperature on benchtop. (E) Ending CD spectrum of PGK that was first denatured in 6 M GdmCl for 24 h, then diluted 100-fold and incubated for 1 h to refold, then ramping upto 95°C at 1 °C/min, and then held at 95°C for 24 h.

**Fig. S3.**
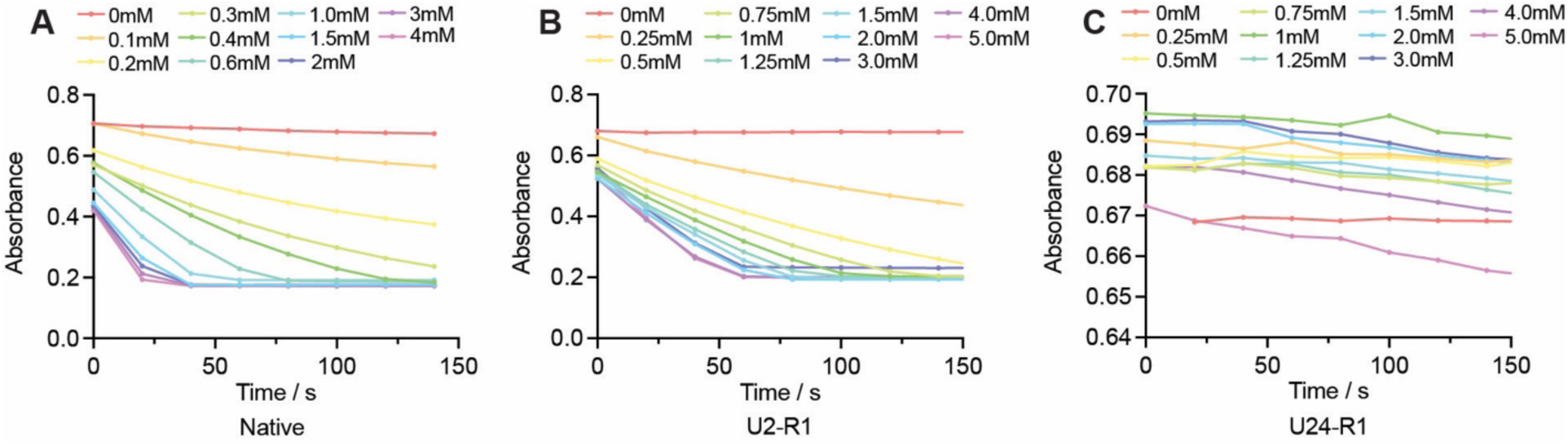
Representative Raw Enzyme Kinetics Data for PGK Variants. (A-C) Representative absorbance traces showing the decrease in NADH signal over time during enzyme assays for (A) native PGK; (B) PGK unfolded for 2 h in 6 M GdmCl, diluted 100-fold and incubated for 1 h (U2-R1); (C) PGK unfolded for 24 h in 6 M GdmCl, diluted 100-fold and incubated for 1 h (U24-R1).

**Fig. S4.**
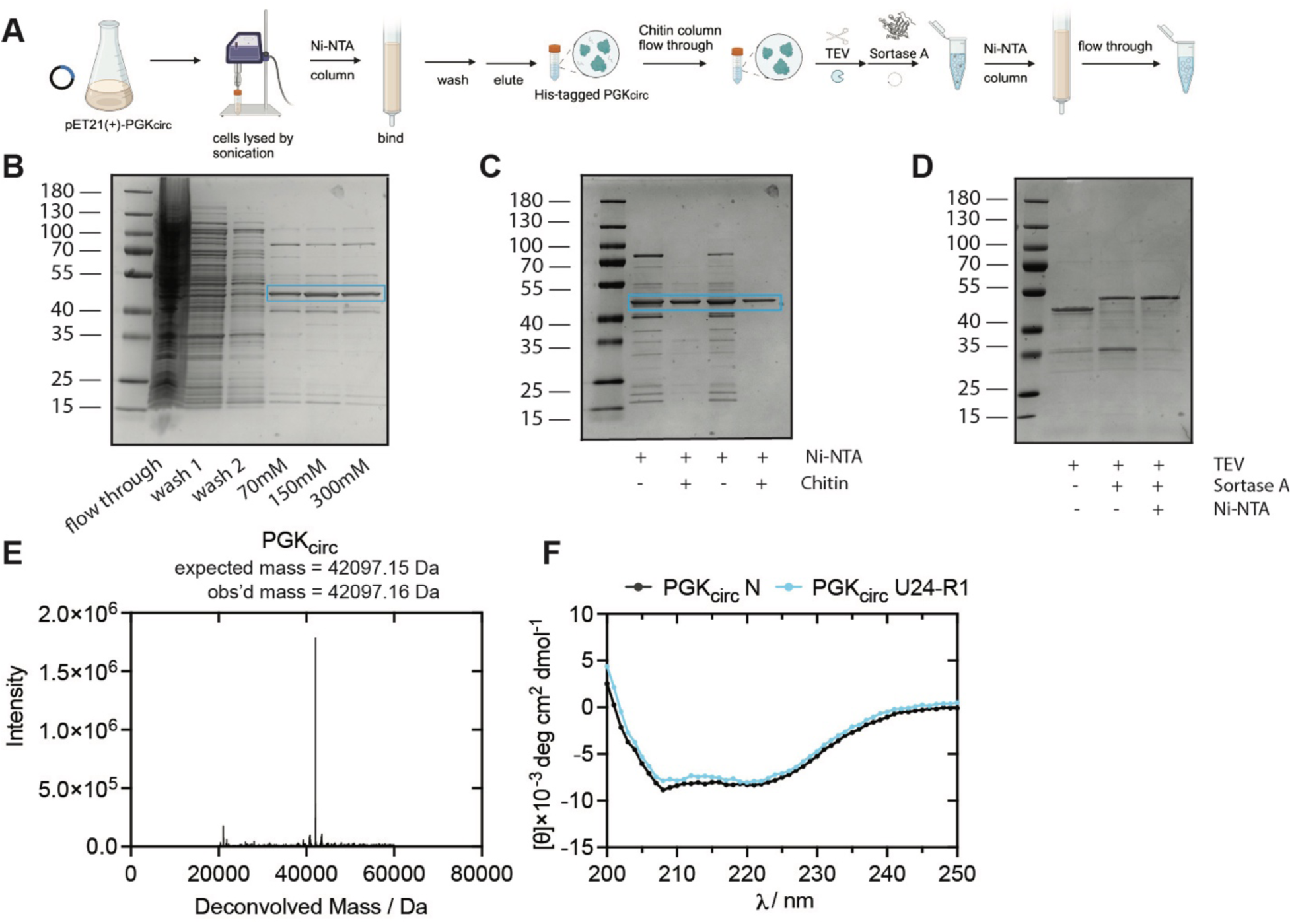
Purification and Characterization of Circularized PGK (PGK_circ_). (A) Scheme illustrating the procedure to purify PGK modified at its N- and C-termini with a circularizable motif using Sortase A. (B) SDS-PAGE stained with Coomassie showing impure pre-PGK_circ_ in the elution fraction from Ni-NTA affinity chromatography. (C) SDS-PAGE stained with Coomassie showing nearly pure pre-PGK_circ_ following depletion through chitin column. Image shows two replicates with and without chitin column treatment. (D) SDS-PAGE stained with Coomassie showing the progression from pre-PGK_circ_ to purified PGK_circ_ by treating with TEV, Sortase A, and depleting enzymes through reverse Ni-NTA chromatography. Circularized proteins are expected to run with greater apparent MW by SDS-PAGE. (E) Max-entropy deconvolved intact mass spectrum confirming the molecular weight of the purified PGK_circ_. (F) Far-UV circular dichroism (CD) spectra of native PGK_circ_ and chemically unfolded-refolded PGK_circ_. Spectra were acquired on 0.1 mg/ml PGK_circ_ in 20 mM HEPES-NaOH pH 7.4, 100 mM NaCl, 2 mm MgCl_2_, 0.06 M GdmCl, 0.1 % (v/v) glycerol, 0.1 mg/ml PGK.

**Fig S5.**
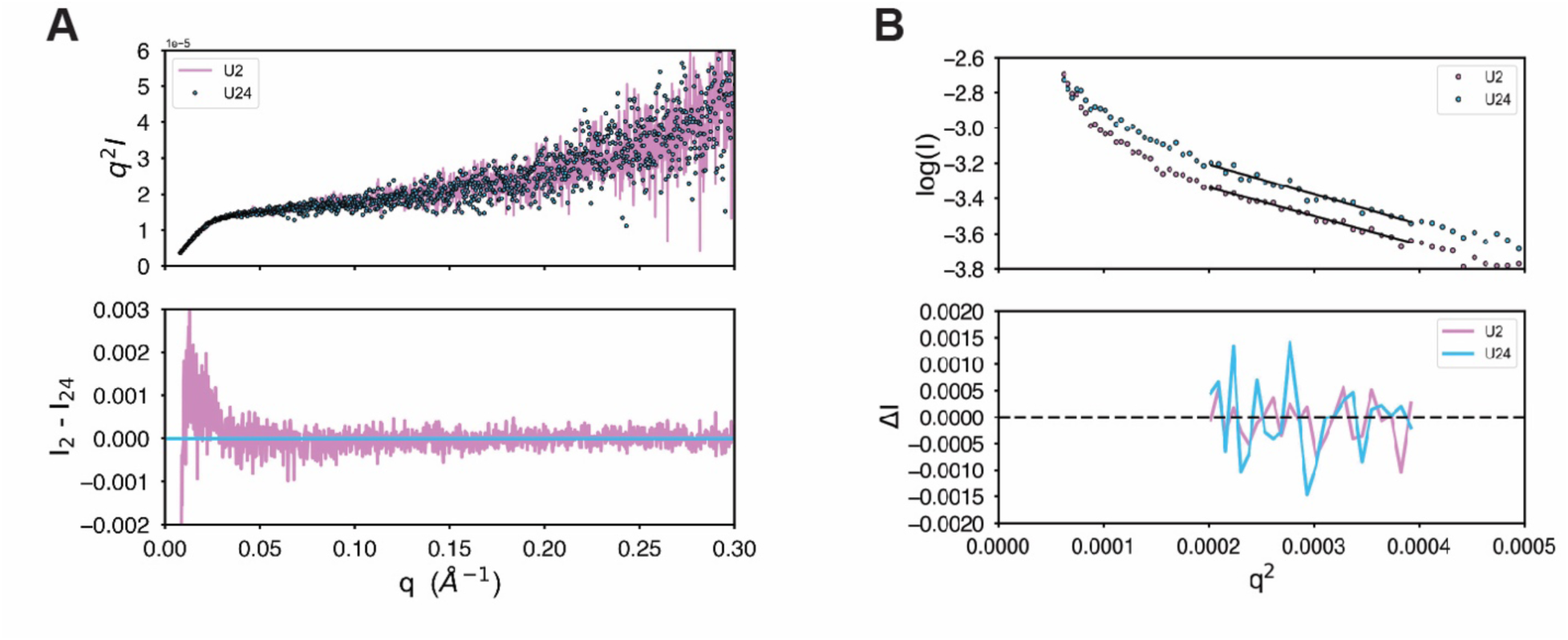
Comparison of SAXS Scattering Profiles for the Unfolded PGK Ensembles. (A)The Kratky plots (top) overlay with a goodness-of-fit *c*2 = 1.06, indicating differences are mostly within experimental noise limits. The plot does not return to baseline at wide angle, a characteristic of unfolded systems. The difference plot (bottom) shows some systematic deviation on very large size scales (small angles) that are typically attributed to unremoved aggregates, X-ray damage, or beam instability. (B) Guinier analysis of U2 and U24 (top). Nonlinearity at smallest angles *q*_2_ < 0.0002 indicates some aggregation or unremoved particulates from the reconstitution process. Black lines show the range of *q* used in the analysis. The difference plot, Δ*I* (bottom), shows no systematic deviations in that range.

**Figure S6.**
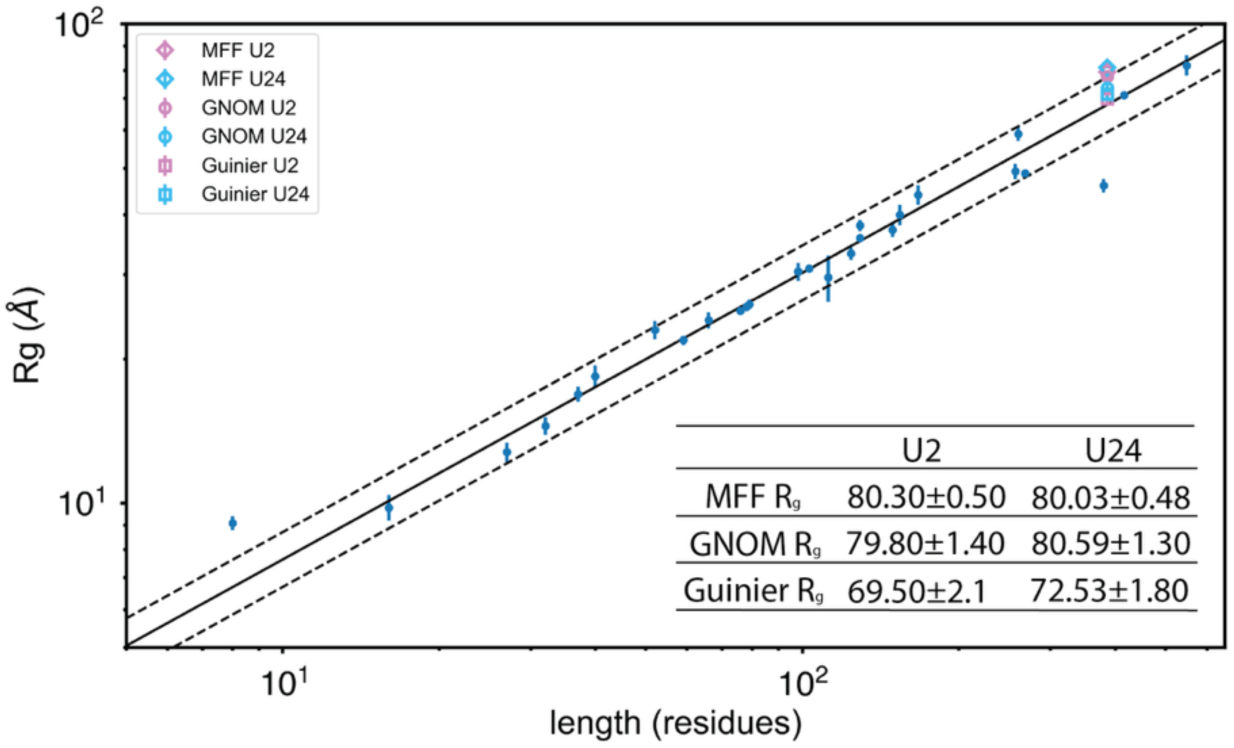
Radius of Gyration, R_g_, of Unfolded PGK. Radius of gyration estimates for U2 and U24 unfolded PGKs as derived by three methods: Molecular form factor (MFF), indirect Fourier transform (GNOM), and Guinier analysis (*85, 86*). The power-law relationship, *R_g_ = R_0_N^v^*, where N = number of residues, *R_0_*= 1.927, and v = 0.598, is shown with the original 28 crosslink-free, prosthetic-group-free, chemically denatured polypeptides tabulated by Kohn et al (*87*). Though at the upper end of the studied protein chain lengths, the U2 and U24 unfolded PGKs both fall within the range expected for an excluded-volume random coil.

**Figure S7.**
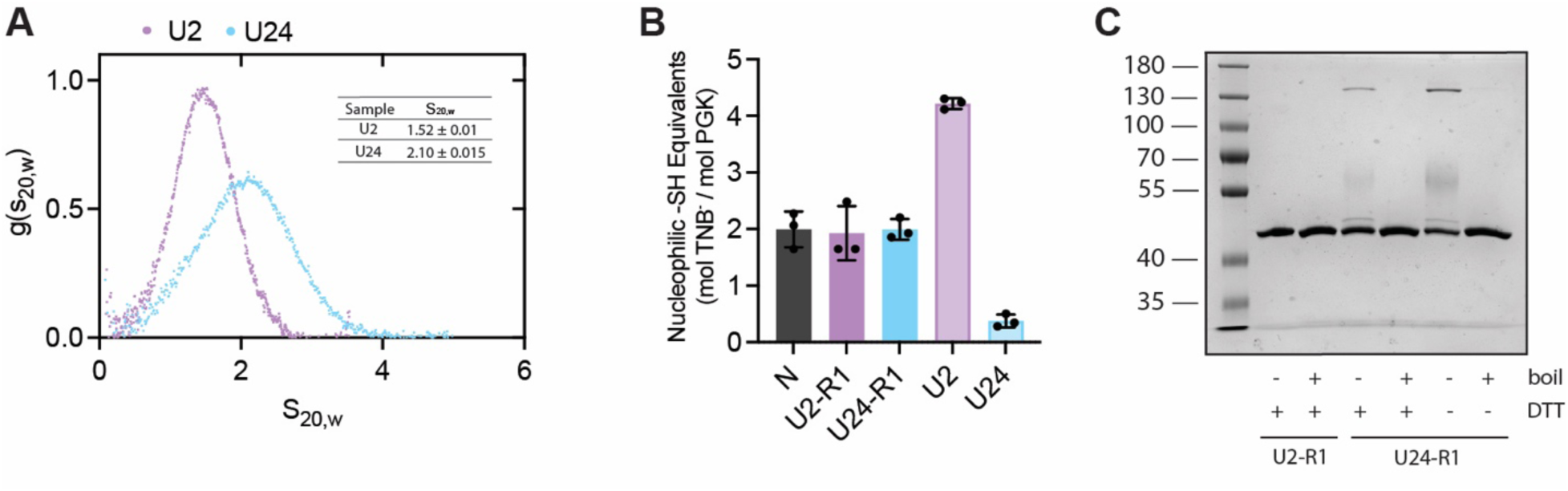
Disulfides can form between unfolded PGK molecules in 8 M urea. (A) Representative sedimentation velocity distributions for unfolded forms of PGK incubated in 8 M urea for the indicated periods of time. Inset table shows mean ± std. dev. for sedimentation velocity normalized to 20°C in pure water (S_20,w_) from technical replicates (*n* ≥ 3). (B) Ellman’s assay to measure the number of nucleophilic thiol (-SH) groups in unfolded and unfolded-refolded PGK samples. Dots indicate replicates (n = 3), bar heights represent means, error bars represent std. devs. (C) Chemically unfolded-refolded forms of PGK, unfolded with (10 mM) or without DTT, were incubated in 0.5 % SDS, boiled (or not), resolved by Tris-Glycine SDS-PAGE, and visualized with Coomassie stain.

**Table S1.** Compilation of sedimentation velocity values from all analytical ultracentrifugation runs.

| PGK form | Buffer | Rep. | Peak Position |  | Gaussian Fit |  |  |  |
| --- | --- | --- | --- | --- | --- | --- | --- | --- |
| | | | $S_{20,w}$ | Error | $S_{20,w}$ | Error | MW | Error |
| Native | A | 1 | 3.202 | 0.021 | 3.162 | 0.002 | 37.90 | 0.26 |
| Native | A | 2 | 3.086 | 0.02 | 3.115 | 0.001 | 40.95 | 0.16 |
| Native | A | 3 | 3.054 | 0.02 | 3.098 | 0.003 | 33.17 | 0.28 |
| U2-R1 | A | 1 | 3.113 | 0.03 | 3.127 | 0.003 | 38.77 | 0.41 |
| U2-R1 | A | 2 | 3.012 | 0.02 | 3.132 | 0.002 | 36.7 | 0.25 |
| U24-R1 | A | 1 | 3.112 | 0.03 | 3.138 | 0.002 | 42.70 | 0.29 |
| U24-R1 | A | 2 | 3.096 | 0.03 | 3.118 | 0.002 | 40.08 | 0.33 |
| U24-R1 | A | 3 | 3.028 | 0.02 | 3.090 | 0.002 | 35.3 | 0.22 |
| U2 | B | 1 | 1.516 | 0.010 | 1.531 | 0.001 | 46.12 | 0.18 |
| U2 | B | 2 | 1.480 | 0.008 | 1.520 | 0.001 | 46.98 | 0.15 |
| U2 | B | 3 | 1.470 | 0.008 | 1.506 | 0.001 | 47.02 | 0.17 |
| U24 | B | 1 | 2.023 | 0.03 | 2.081 | 0.002 | 41.47 | 0.33 |
| U24 | B | 2 | 2.097 | 0.03 | 2.109 | 0.001 | 41.37 | 0.23 |
| U24 | B | 3 | 2.023 | 0.03 | 2.083 | 0.002 | 40.92 | 0.31 |
| U24 | B | 4 | 2.074 | 0.03 | 2.108 | 0.001 | 41.64 | 0.23 |
| U2 | C | 1 | 1.491 | 0.03 | 1.46 | 0.002 | 34.7 | 0.43 |
| U24 | C | 1 | 1.496 | 0.05 | 1.438 | 0.003 | 30.6 | 0.5 |

All samples were measured with 0.1 mg/ml PGK in 25°C at 235 nm. $S_{20,w}$ values (sedimentation coefficients normalized to 20°C in pure water) were tabulated in one of two ways: either by extracting the peak position from the distributions or by fitting to a Gaussian model (81). Buffer A is 20 mM HEPES-NaOH pH 7.4, 100mM NaCl, 2 mM MgCl<sub>2</sub>, 0.06 M GdmCl, 0.1% glycerol. Buffer B is 20 mM HEPES-NaOH pH 7.4, 100mM NaCl, 2 mM MgCl<sub>2</sub>, 8 M urea. Buffer C is 20 mM HEPES-NaOH pH 7.4, 100mM NaCl, 2 mM MgCl<sub>2</sub>, 8 M urea, 1 mM DTT.

**Table S2.**
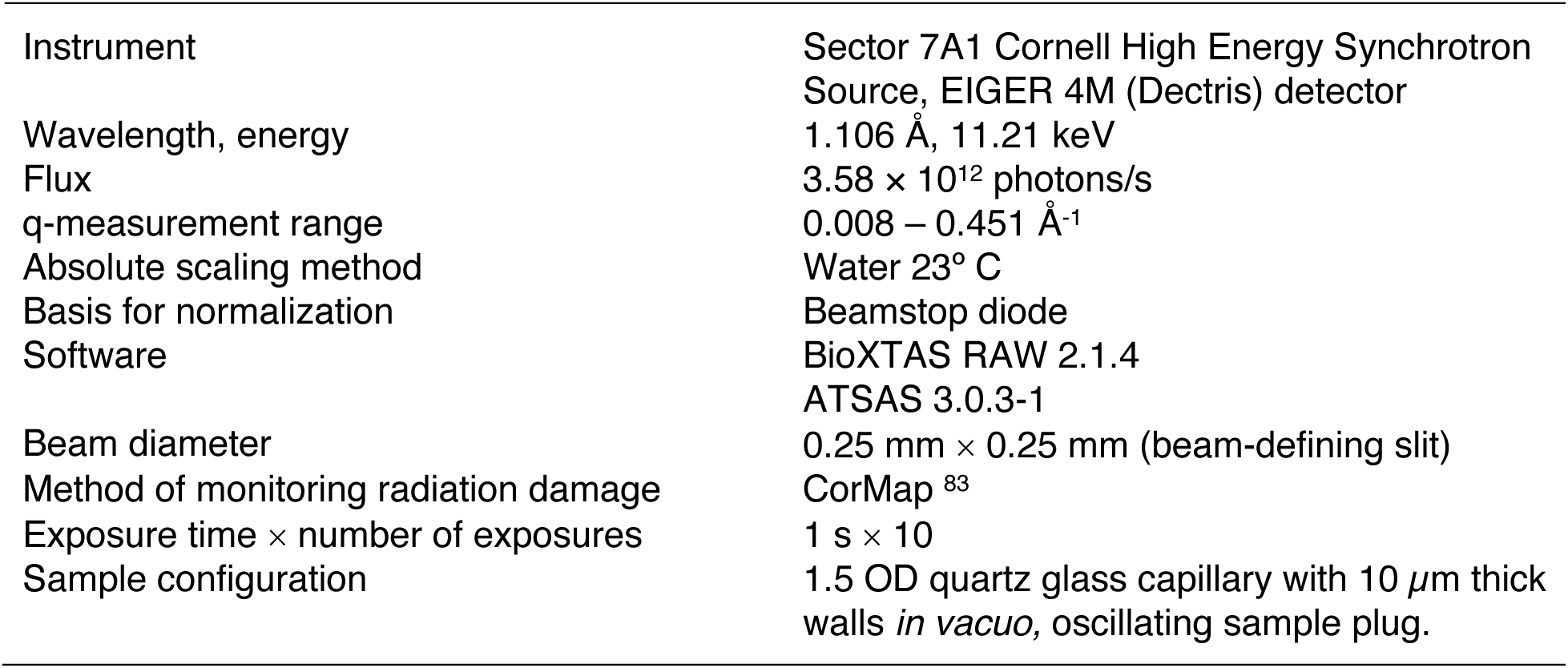
SAXS Acquisition Parameters.

**Other Supplementary Material for this manuscript includes the following:**

Data S1–S3.

---

**Data S1.** DataS1-CD_RawData.xlsx

Contains all the CD spectra shown in the manuscript and supplemental materials

**Data S2.** DataS2-AUC_RawData.xlsx

Contains all the AUC sedimentation distributions shown in the manuscript and supplemental materials

**Data S3.** DataS3-Activity_RawData.xlsx

Contains all the initial rates from enzyme activity assays shown in the manuscript

## Notes

### Competing Interest Statement

The authors have declared no competing interest.

